# Bonds, a neighbourhood field and missing hydrogens: a transferable electron-density model of proteins

**DOI:** 10.64898/2026.11.26.769548

**Authors:** Yongli Shan, Humayun Sharif, Hanning Chen

**Author notes:** Corresponding authors: Hanning Chen,; Yongli Shan,.

## Abstract

A compound that has not been made has no diffraction data to refine against. A model of electron density that is to be used in drug design must therefore be transferable: the same numbers for every protein and every ligand, nothing fitted per structure. The independent-atom model (IAM) is transferable, but it places no density in bonds and lets no atom respond to its surroundings. We set out to add both, explicitly and with few parameters, and tested each step on every Protein Data Bank X-ray entry with structure factors at 1.0 Angstrom or better (2,006 entries; 650 proteins never seen in training). Four things were learnt on the way. The benchmark itself had to be corrected first: in models deposited without hydrogen atoms, placing riding hydrogens lowers the R of the IAM by 0.0093 in 315 of 316 proteins, and about four fifths of what a transferable multipole model appeared to gain on them was the missing hydrogens. With hydrogens placed on every model, a layer that names each bond and contact, gives it one charge and one width and takes that charge from the valence shells of its atoms (the Complete Electron-cloud Layer, CEL) lowers R in 611 of 650 proteins, but by only 0.0007. Most of what the IAM leaves out is centred on the atoms and set by their surroundings. A neighbourhood field of 3,055 parameters, computed by matrix products from the types and distances of all atoms within 8 Angstrom, lowers R three times more than the bond layer, and the two together (3,214 parameters) are better than the multipole (4,768) in 545 of 650 proteins. And a bond has to be named by its chemical environment: with one class per pair of elements the C-N bond came out negative; with 25 classes defined beforehand the peptide bond is positive, the carbon skeleton carries 0.04 to 0.12 electrons per bond in a chemical order, and the layer is better on 602 of 650 unseen proteins. Read inversely, the bonds of the main chain stand out of the density in more than four proteins of five, and the polar bonds of ligands, which read below empty points in the density left by independent atoms, are restored by the field predicted from the surroundings, with its factor calibrated on ligand bonds against the per-atom solution. A per-atom solution of the same radial form lowers R by 0.017 on the reflections it is fitted to, but by about 0.002 on reflections held out of the fit, which is the size of what the field predicted from distances already delivers: the spherical form is close to what the data support, and the next gain has to come from direction. The bonds respond to their surroundings as well, but as a secondary effect: freed bond by bond and tested on held-out reflections, the charge of a bond changes by about 0.06 electron and gives one tenth of the gain of the atoms, and no shift of the bond charge is supported by the data.

**Synopsis:** A transferable electron-density model for proteins, made of a named bond layer that conserves electrons and a neighbourhood field computed from interatomic distances, is benchmarked on 650 unseen sub-Angstrom proteins; with hydrogens placed on every model it is better than a transferable multipole model in four proteins out of five, and most of the multipole’s apparent gain on models deposited without hydrogens is shown to be the hydrogens.

## 1. Introduction

### 1.1 A transferable density model for structure-based design

Structure-based drug discovery reasons about a compound in a protein pocket before the compound is synthesised. What decides binding is the electron density of the two molecules where they meet: the bonds inside each, and the contacts between them. A model of that density is useful for design only under three conditions. It must be transferable, because a designed compound has no data of its own. It must be explicit, each number naming a bond, a contact or an environment, so that a prediction can be traced to chemistry and checked. And it must respond to the surroundings of an atom, because binding is exactly a change of surroundings. This paper builds such a model and tests it where it can be tested against measurement: in sub-Angstrom protein crystals, where the density beyond independent atoms is in the data. No binding calculation is reported here; the paper establishes the density model that such a calculation needs.

### 1.2 Existing models of electron density

Routine refinement uses the independent-atom model (IAM): one spherical, neutral scattering factor per element. It is transferable and it is wrong in a known way: no density in bonds, no response of an atom to its neighbours. Multipolar refinement (Hansen & Coppens, 1978) and its spherical subset, the kappa formalism (Coppens et al., 1979), expand each atom in atom-centred functions fitted on the structure. Transferable aspherical-atom libraries move multipole parameters from small molecules to proteins (Jelsch et al., 2000; Pröpper et al., 2013), with one set of populations for each atom type in its bonded environment. Interatomic scatterers place one Gaussian on each bond and refine its charge on the structure (Afonine et al., 2007, 2012). These models were demonstrated on single structures or on a handful. None was tested across the archive for what a design model must do: predict reflections it has not seen, in proteins it has not seen, with nothing fitted on the entry.

### 1.3 This work

We build the model in two explicit parts and test it on the whole sub-Angstrom archive. The first part is a bond layer in which every covalent bond and every named contact carries one charge and one width, with the charge taken from the valence shells of the atoms that form it (section 2.2); its covalent classes are defined beforehand by chemical environment (section 2.3). The second part is a neighbourhood field that computes a spherical correction of every atom from the types and distances of all atoms around it (section 2.4). Both are compared with a transferable multipole model trained on the same data (section 2.5), on a benchmark in which every model carries hydrogens placed by one rule (section 3). The results (section 4) establish four points.

1. Bonds. A layer of 159 named numbers, conserving electrons, lowers R on held-out reflections in 611 of 650 unseen proteins (sections 4.2 and 4.3).
2. The neighbourhood field. A term of 3,055 parameters computed from interatomic distances lowers R three times more than the bond layer, and with the bond layer is better than the transferable multipole in 545 of 650 proteins (section 4.6).
3. Hydrogens. The benchmark of any such model depends on whether the deposited model contains hydrogen atoms. Where it does not, most of what a transferable multipole model gains is the density of the missing hydrogens (section 4.1). All comparisons are therefore made with hydrogens placed on every model by one rule.
4. Bonds resolved by chemical environment. A class of bonds defined by its two elements mixes bonds that are different. Defined beforehand by chemical environment, the classes separate, the charges become readable one by one, and the gain of the bond layer on unseen proteins rises by 60 per cent (section 4.3).

**Table 1.** The results at a glance (hydrogens placed on every model; 650 proteins never seen in training; better : worse counts proteins; dR is the median over proteins of the change of R).

| test | result | number | where |
|---|---|---|---|
| Effect of placed hydrogens | IAM with riding hydrogens against IAM as deposited, models deposited without H | 315 : 1, dR -0.0093 | section 4.1 |
|  | multipole against IAM on those proteins, as deposited and with H | -0.0091 and -0.0019 | section 4.1 |
| Gain over the IAM | multipole (4,768 parameters) | 589 : 61, -0.0018 | section 4.2 |
|  | CEL, electrons conserved (159) | 611 : 39, -0.0007 | section 4.3 |
|  | CEL + neighbourhood field (159 + 3,055) | 595 : 55, -0.0029 | section 4.6 |
| Bond classes by chemical environment | bond layer, classes by chemical environment and one kappa per element (159) against classes by element pair (104) | 602 : 48, -0.0003 | section 4.3 |
|  | the complete model on the one bond layer against the other | 480 : 170, -0.0003 | section 4.6 |
| Comparison with the multipole | CEL + field against multipole | 545 : 105, -0.0012 | section 4.6 |
|  | the same, field with 9,221 parameters | 557 : 93, -0.0015 | section 4.6 |
|  | the same, field trained on the multipole's training set only | 414 : 236, -0.0003 (9,221 parameters: 476 : 174, -0.0008) | section 4.7 |
| Field on a multipole base | field on multipole + CEL layer, hydrogens placed (647 proteins) | 570 : 77, -0.0018 | section 4.7 |
| Upper bound of the spherical form | per-atom solution fitted per entry, R on the reflections of the fit | 650 : 0, -0.0173 | section 4.8 |
|  | the same, R on reflections held out of the fit (48 entries of the training tier) | 36 : 12, -0.0022 | section 4.8 |
| Environment dependence of bond charges | charge of every bond freed, R on reflections held out of the fit (109 entries of the training tier) | 83 : 26, -0.0002 | section 4.9 |
|  | the same, dipoles of every bond freed | 49 : 60, +0.0001 | section 4.9 |
| Inverse read-out of bonds | main-chain CA-C bond above matched empty points | 568 : 60, AUC 0.62 | section 4.10 |
|  | polar bonds of ligands (C-O), density against density minus predicted field (per-atom solution) | AUC 0.417 and 0.510 (0.500) | section 4.10 |

## 2. Model

The model adds two explicit parts to the independent-atom model: a bond layer that conserves electrons and a neighbourhood field computed from the surroundings of every atom. The need for the first is visible in the difference map of the IAM, which, averaged over the peptide units of a crystal, shows structured density in the peptide plane (figure 1).

**Figure 1.**
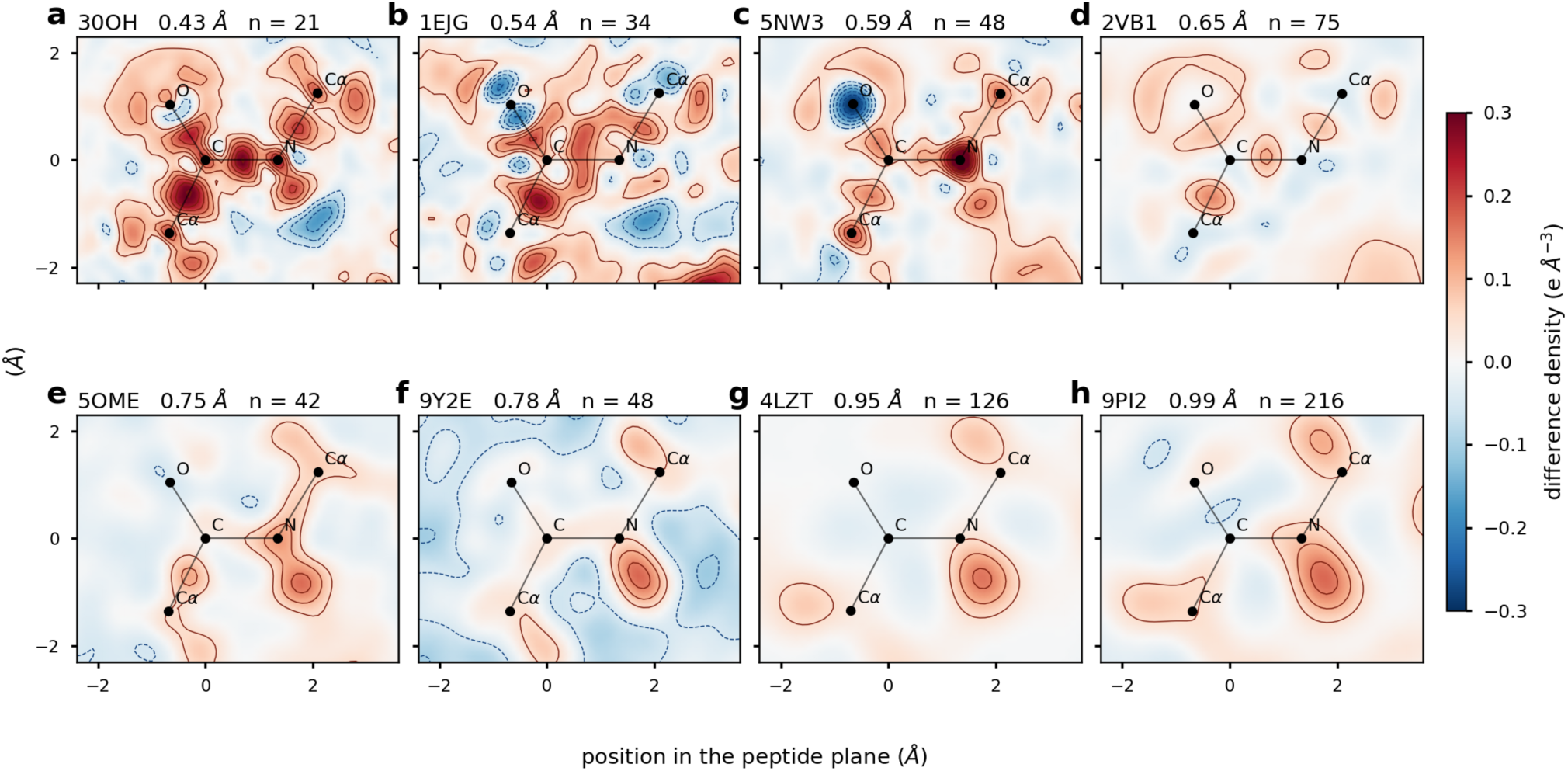
Residual density of the independent-atom model in the peptide plane. Mean difference density (e A^-3) over the n peptide units of each of eight crystals, in order of resolution; each title gives the PDB entry, d_min and n. Solid contours are positive and dashed contours negative, in steps of 0.05 e A^-3; the colour scale is common to the eight panels. Dots and lines mark CA, C, O, N and the following CA. Both axes are positions in the peptide plane in A (`fig1_peptide_maps`).

### 2.1 Independent-atom base

The base is the IAM with the deposited coordinates, occupancies and displacement parameters. Scale, a two-parameter bulk solvent and one overall B are fitted on each entry, for every model alike. Nothing else is fitted on an entry.

### 2.2 Electron-conserving bond layer

Every covalent bond type present in the training set, and thirteen non-covalent interaction classes with explicit geometric definitions (three hydrogen-bond classes, salt bridge, halogen bond, parallel and T-shaped aromatic stacking, cation-pi, chalcogen, n-to-pi* carbonyl, C-H-pi, sulfur-pi and amide-pi), is a class t. Each class has one charge *q_t_* and one width *w_t_*. A Gaussian of that charge and width is placed at the bond midpoint or at the contact point of every member of the class:

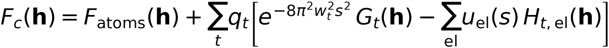

*G_t_* is the sum over the sites of the class (with their occupancy, the mean displacement of their two partners and the symmetry operators), as in the additive layer of the earlier version. The second term is what makes the layer conserving: the charge placed on a site is removed from the atoms that form it. *H_t, el_* sums over the donor atoms of element el, each with its own displacement and with the share it gives: both atoms of a covalent bond give half; the two partners of a contact give half each, and an aromatic ring divides its share equally among its atoms. *u_el_*(*S*) is the normalised form factor of the valence shell of the element, so that the charge is taken from the valence shell and the core is untouched. At *q* = 0 the model is the IAM. An atom’s cloud is thus one whole: the part that stays on the atom, and the part that the atom gives to its bonds and contacts. No atom gives more than its valence shell holds in any of the 2,006 entries.

Variants that also let the inner shell give a share, or let its displacement differ, were trained and change nothing (section 4.2): we find no evidence that the inner shell responds to bonding.

### 2.3 Bond classes by chemical environment

Two definitions of the covalent classes are used, and they are kept apart in what follows. In the first a class is a pair of elements (C-C, C-N, C-O and so on; 39 pairs occur, most of them in ligands and cofactors); with the thirteen contact classes this layer has 52 classes and 104 parameters. It is the control in what follows (section 4.2). In the second the covalent classes are defined beforehand by chemical environment, not by the two elements alone. For the twenty standard residues the class of a bond follows from the names of its two atoms: carbon-carbon bonds are main-chain CA-C, aromatic, sp3 to aromatic, sp3 to a side-chain carbonyl or carboxyl carbon, and sp3 to sp3; carbon-nitrogen bonds are the peptide bond, N-CA, the proline N-CD bond, aromatic, side-chain amide, guanidinium and amine; carbon-oxygen bonds are main-chain carbonyl, side-chain amide carbonyl, carboxylate, hydroxyl and phenol; carbon-sulfur bonds are thioether and cysteine; bonds to hydrogen are classed by the atom that carries the hydrogen (C-H on CA, aromatic, sp3; N-H main chain, charged side chain, other side chain). These are 25 classes. A bond that the rules do not place, in a ligand, a modified residue or at a metal, keeps the class of its two elements, so that no bond is left out. The rules are fixed before any fit. With this definition each element also has one contraction coefficient kappa for its valence shell, with the valence population held at that of the neutral atom,

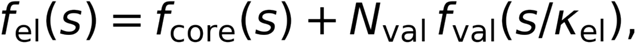

so that the electron count is again unchanged. This layer has 77 classes and 159 parameters (section 4.3); it is the bond layer of the model.

### 2.4 Neighbourhood field

The second part of the model is a correction on every atom, spherical to first order, different for every individual atom, and computed from the positions and kinds of the atoms around it; section 4.5 shows that the density left by the bond layer has these properties. The neighbourhood field is the smallest model we could write that meets them.

After the bond layer, each atom a carries a spherical correction of its cloud, a radial function *W_a_*(*r*) on 17 nodes from 0 to 1.6 Angstrom (zero at 1.7 Angstrom), entering the structure factor through its three-dimensional Fourier transform:

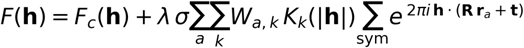

*W_a_* is not fitted on the entry. It is computed from the matrix of distances between all atoms of the crystal within 8 Angstrom of a, bonded or not, and their types. With S the sparse matrix that distributes each pair of atoms over radial basis functions of their distance,

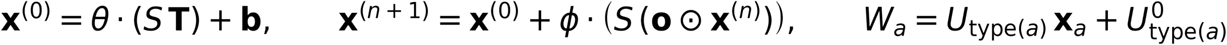

where T is the one-hot matrix of atom classes, o the occupancies, and theta, *b*, phi, *U*, *U*^0^ are the parameters: 3,055 with one channel, 9,221 with three channels and the logarithm of the atomic B as an input. The first line is the direct influence of every neighbour on the atom; the iteration lets the state of every neighbour, itself set by its own neighbours, act in turn, so that the field at an atom integrates the network of all atoms and not only the pairs it belongs to. All operations are sparse matrix products. lambda is a single number for the whole benchmark. sigma is the standard deviation of the difference map of the entry and sets the unit.

The field is learnt from the per-atom solution of section 4.8: on each training entry the radial function of every atom is solved from the difference map left by the bond layer, and the parameters are fitted to reproduce those functions from distances and types. It is then evaluated only on proteins that were not used for that fit.

### 2.5 Transferable multipole reference

The reference is a Hansen-Coppens multipole model (l up to 3) in our own implementation, with one set of valence population, contraction and multipole populations for each atom type defined by element and bonded environment (287 types, 4,768 parameters), trained on the same training entries and transferred without a fit, exactly as the CEL. We also report the CEL as an additional layer on that multipole model.

## 3. Benchmark

Every model is judged by whether it lowers R on reflections and in proteins that it has not seen, with hydrogens placed on every model by one rule.

### 3.1 Data sets and evaluation by protein

Training set T1: 111 entries (two folds); test set T2: the 2,006 entries of the tier, 1,996 computed. A T2 entry is evaluated with parameters trained on the other fold. Proteins are the unit: the entries are grouped into 726 proteins, every protein with an entry in T1 is removed, and each of the remaining 650 proteins counts once, by the median over its entries. Reported are the number of proteins where a model has the lower R against the number where it has the higher, and the median over proteins of the difference dR. Collapsing by batch before collapsing by protein, or counting entries, does not change any conclusion of this paper (section 4.7); it did change them before the hydrogens were placed.

### 3.2 Riding hydrogens on every model

Of the 1,996 entries, 1,299 were deposited with hydrogen atoms and 697 without; by protein, 297 with, 316 without, 37 mixed. A model without hydrogens is missing about one electron for every hydrogen, at a known place. Any model with free atom-centred populations, trained on such entries, can take those electrons up.

We removed the deposited hydrogens from every entry and placed all hydrogens again by one rule: riding positions from the CCP4 monomer library (Vagin et al., 2004) with gemmi (Wojdyr, 2022), electron-cloud X-H distances, none on water molecules, occupancy and alternative-conformer label of the parent atom, B of the hydrogen 1.2 times the equivalent B of its parent. 4,662,006 hydrogens were placed on 5,414,890 heavy atoms; no entry was lost; monomers without a library entry (the most frequent in 47 and 33 entries) received none. On entries deposited with hydrogens the rule reproduces the R of the deposited model (four test entries: 0.1109 against 0.1106, 0.1245 against 0.1240, 0.1642 against 0.1640, 0.1555 against 0.1546), and electron-cloud distances give a lower R than internuclear ones in every case tested.

## 4. Results

### 4.1 Effect of missing hydrogens on the benchmark

Missing hydrogens dominate the apparent gain of an atom-centred model on deposited coordinates (table 2).

**Table 2.** The effect of hydrogens (650 proteins; better : worse, median dR).

| **comparison** | **deposited without H (316)** | **deposited with H (297)** | **all 650** |
|---|---|---|---|
| IAM with hydrogens placed against IAM as deposited | 315 : 1, -0.0093 | 125 : 172, +0.0001 | 476 : 174, -0.0041 |
| multipole against IAM, models as deposited | 310 : 6, -0.0091 | 231 : 66, -0.0012 | 577 : 73, -0.0043 |
| multipole against IAM, hydrogens placed | 297 : 19, -0.0019 | 257 : 40, -0.0017 | 589 : 61, -0.0018 |
| CEL (conserving) against IAM, models as deposited | 292 : 24, -0.0005 | 254 : 43, -0.0004 | 581 : 69, -0.0005 |
| CEL (conserving) against IAM, hydrogens placed | 285 : 31, -0.0005 | 251 : 46, -0.0004 | 572 : 78, -0.0005 |

Three things follow (figure 2). Placing hydrogens on a model deposited without them lowers the R of the IAM by 0.0093 in 315 of 316 proteins, twice what the multipole model gains on the whole set. On those proteins the multipole gains 0.0091 as deposited and 0.0019 once the hydrogens are there: about four fifths of its gain was the hydrogens. And with hydrogens in place the two classes of proteins behave alike (0.0019 and 0.0017), where before they differed by a factor of eight. Whether the deposited model contains hydrogens was the strongest predictor of the multipole gain among all properties of an entry that we recorded (rank correlation 0.83; resolution, mean B, completeness and the coverage of atom types all below 0.3).

**Figure 2.**
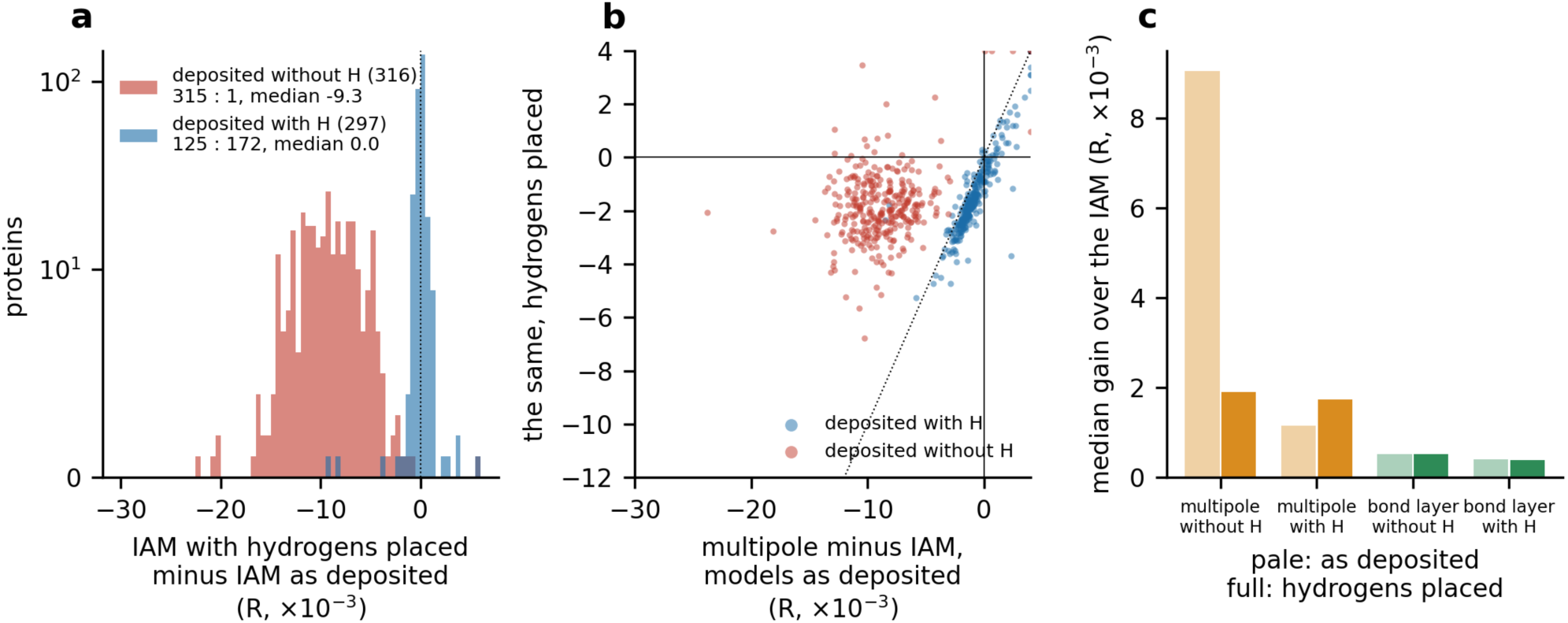
Hydrogens decide what the benchmark measures (section 4.1). Each of the 650 proteins once. (a) Change of the R of the IAM when hydrogens are placed, for proteins deposited without hydrogens (316) and with hydrogens (297); the vertical axis is linear to 10 and logarithmic above; values are clipped at -30 and +6. (b) Gain of the multipole over the IAM with hydrogens placed against the same gain on the models as deposited, one point per protein; dotted line: equal; values beyond the axes are drawn at the edge. (c) Median gain over the IAM of the multipole and of the bond layer by element pair, by whether the model was deposited with hydrogens; pale: as deposited, full: hydrogens placed (`fig14_hydrogens`).

The bond layer does not show the effect: its gain is the same with and without hydrogens. A layer whose charges sit on bonds and are taken from the atoms cannot stand in for a missing atom.

The consequence for method comparison is general. A benchmark of a density model on deposited coordinates mixes two things, the description of bonding and the completion of the model, and the mixture depends on the refinement habits of the depositors. All comparisons below are therefore made with hydrogens placed on every entry (section 3.2), and all parameters of every model were trained again on the training set with hydrogens.

### 4.2 Electron-conserving bond layer

With hydrogens placed, the electron-conserving bond layer lowers R in most proteins, by a small amount (table 3).

**Table 3.** Bond layers with hydrogens placed (650 proteins).

| model | parameters | against IAM |
|---|---|---|
| CEL, additive (charge added on the bonds, atoms unchanged) | 104 | 624 : 26, -0.0011 |
| CEL, conserving (charge taken from the valence shells) | 104 | 572 : 78, -0.0005 |
| multipole | 4,768 | 589 : 61, -0.0018 |
| CEL as a layer on the multipole, against the multipole | 92 | 536 : 111, -0.0002 (647 proteins) |

Conservation of electrons is the physical statement of the model and is kept in everything below. Its gain is small and consistent. The additive layer gains twice as much with the same number of parameters, and the reason is visible in its charges: without the constraint, classes take up charge that is not theirs (the carbonyl n-to-pi* class runs to its bound of +3 electrons, the C-H class to -1.8); with the constraint the charges fall into a range that can be read (C-C +0.06 to +0.07 electrons at a width of 0.16 A, C-S +0.04 to +0.07, C-H -0.09 to -0.11). One class stands out: C-N is negative and broad in both folds (-0.23 to -0.27 electrons, width 0.7 A). section 4.3 traces it. Two contact classes sit at the upper bound of the width. Letting the inner shell give a share of the charge (four more parameters) or giving it its own displacement changes the result in the fifth decimal; we find no response of the inner shell to bonding.

A hundred numbers on bonds recover about a quarter of what the multipole recovers, so most of what a transferable model can add to the IAM is not at the midpoints of bonds (section 4.4).

### 4.3 Bond classes by chemical environment and the C-N charge

The negative, broad charge of the C-N class is not an effect of the hydrogens (it is -0.32 to -0.35 electrons on the models as deposited) and it is absent from the additive layer (+0.03 to +0.05 electrons, width 0.05 to 0.19 A). It appears with the conservation of electrons. Three tests on the training set, each in both folds, say what it is (table 4).

**Table 4.** The C-N class of the conserving layer (charge in electrons / width in A; two folds).

| **test** | **C-N** | **reading** |
|---|---|---|
| conserving layer, classes by element pair | -0.270 / 0.70, -0.231 / 0.72 | reference |
| one more parameter per polar class: electrons moved from the less to the more electronegative atom of the bond | -0.282 / 0.77, -0.237 / 0.80; moved -0.04, -0.04 | not a substitute for charge transfer between the two atoms |
| C-N in three classes: peptide bond | +0.022 / 0.05, +0.017 / 0.05 | a bond |
| N-CA | -0.590 / 0.94, -0.498 / 0.92 | the whole negative signal |
| all other C-N | +0.028 / 0.05, -0.073 / 0.54 | the folds disagree |
| one kappa per element, classes by element pair | +0.030 / 0.05, +0.024 / 0.05 | a bond, once the valence shell of N may contract (kappa 1.009, 1.005) |

A width of 0.9 A covers both atoms of the bond: the negative charge at the N-CA midpoint, returned to the valence shells of N and CA, makes these two atoms sharper. It is an effect centred on the atoms that a class of bonds was carrying because the model offered no other place for it, and N and CA are the atoms whose centres carry 1.3 to 1.9 per cent more electrons than the IAM in the per-atom solution (section 4.4). The two independent tests agree. The class by element pair had put the peptide bond and the N-CA bond, which behave differently, under one number.

The covalent classes were therefore defined beforehand by chemical environment (section 2.3), all families fitted together on the training set, and the layer evaluated on the 650 unseen proteins (table 5).

**Table 5.** The bond layer by the definition of its classes (hydrogens placed; 650 proteins; 1,996 entries).

| **bond layer** | **parameters** | **against IAM** | **against the layer by element pair** |
|---|---|---|---|
| classes by element pair | 104 | 572 : 78, -0.00045 |  |
| classes by element pair and one kappa per element | 109 | 557 : 93, -0.00043 | 295 : 355, +0.00001 |
| classes by chemical environment | 154 | 604 : 46, -0.00065 | 595 : 55, -0.00021 |
| classes by chemical environment and one kappa per element | 159 | 611 : 39, -0.00072 | 602 : 48, -0.00031 |

The separation holds on proteins the parameters have not seen, in nine out of ten, and it raises the gain of the bond layer by 60 per cent. It is the separation of the classes that does it. kappa alone, with the classes by element pair, changes nothing on unseen proteins (295 : 355), although on the training set it turns the C-N charge positive (table 4); with the classes separated it adds a further 0.00008 (460 : 190). The bond layer alone remains well below the multipole (85 : 565); what the definition changes is that the numbers can now be read class by class (table 6, figure 3a), and the gain holds protein by protein (figure 3b). The layer with classes by chemical environment and kappa is the bond layer of the model from here on.

**Figure 3.**
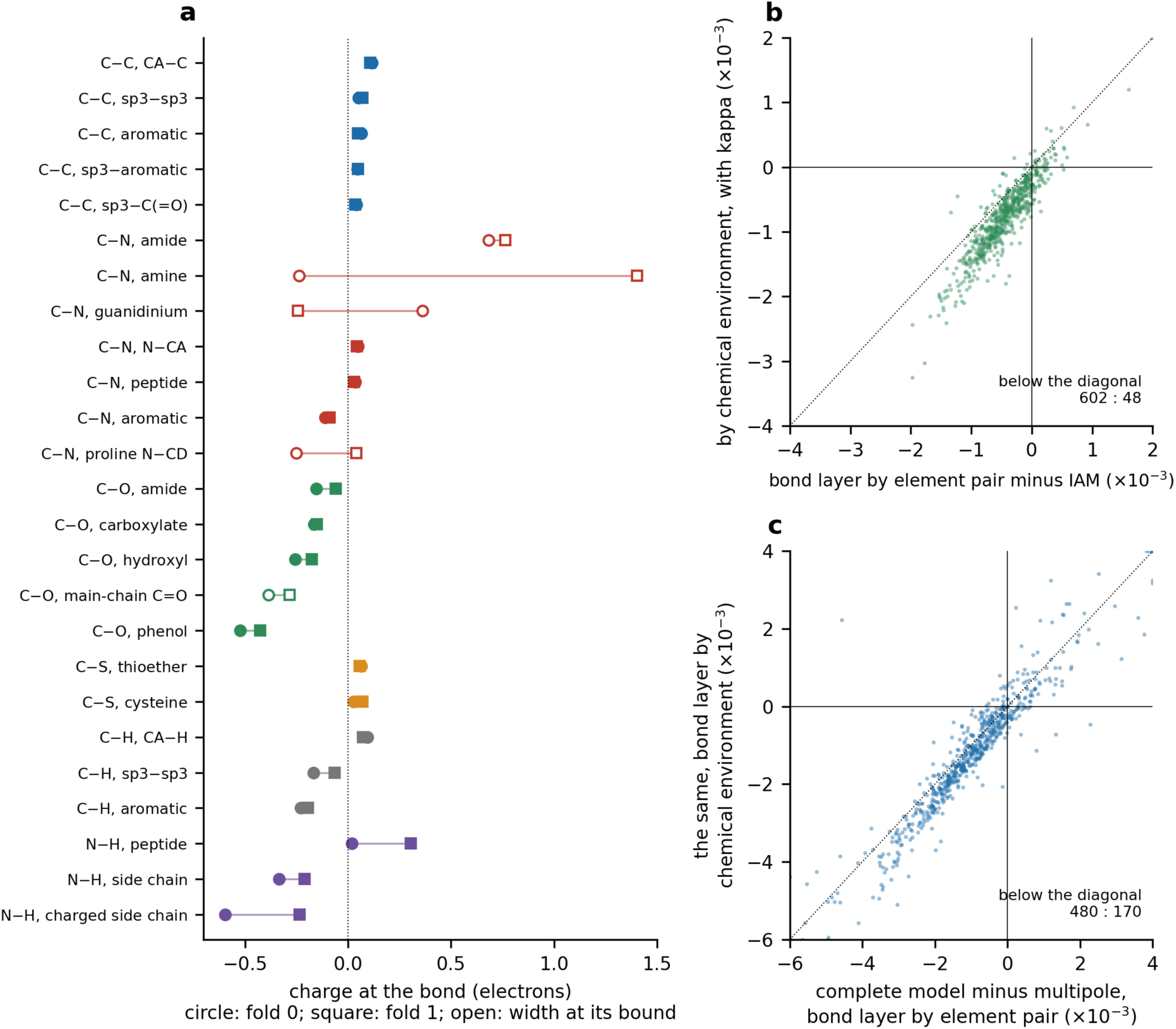
Bonds by chemical environment (section 4.3). (a) Charge of each of the 25 covalent classes of the standard residues in the two folds of the training set (circle and square), grouped by pair of elements and ordered by charge; open symbols: the width is at its upper bound in at least one fold. (b) Gain of the bond layer over the IAM on the 650 unseen proteins, with classes by chemical environment and kappa against classes by element pair; below the dotted diagonal the former is better. (c) The complete model (bond layer + field, one channel) minus the multipole, on the two bond layers. Values beyond the axes are drawn at the edge (`fig16_bond_classes`).

**Table 6.** Charges of the classes that agree in the two folds (classes by chemical environment with kappa; electrons, width in A in brackets).

| **bond** | **fold 0** | **fold 1** |
|---|---|---|
| C-C, main-chain CA-C | +0.117 (0.22) | +0.108 (0.22) |
| C-C, aromatic | +0.066 (0.20) | +0.049 (0.17) |
| C-C, sp3 to sp3 | +0.051 (0.13) | +0.071 (0.20) |
| C-C, sp3 to aromatic | +0.047 (0.10) | +0.048 (0.13) |
| C-C, sp3 to carbonyl or carboxyl carbon | +0.041 (0.07) | +0.035 (0.09) |
| C-N, peptide bond | +0.035 (0.10) | +0.030 (0.07) |
| C-N, N-CA | +0.049 (0.12) | +0.043 (0.14) |
| C-S, thioether | +0.065 (0.05) | +0.056 (0.13) |
| C-S, cysteine | +0.028 (0.12) | +0.070 (0.20) |
| S-S | +0.089 (0.25) | +0.184 (0.19) |
| C-O, carboxylate | -0.163 (0.31) | -0.151 (0.31) |
| C-O, hydroxyl | -0.255 (0.46) | -0.174 (0.42) |
| C-O, phenol | -0.524 (0.45) | -0.425 (0.45) |
| C-H, aromatic | -0.229 (0.48) | -0.195 (0.45) |
| C-H, sp3 | -0.167 (0.48) | -0.065 (0.43) |
| N-H, side chain | -0.334 (0.41) | -0.209 (0.35) |
| N-H, charged side chain | -0.595 (0.34) | -0.234 (0.31) |

The bonds of the carbon skeleton are positive and narrow in every class, with an order that chemistry expects: the main-chain CA-C bond, next to the carbonyl, carries about 0.11 electrons, aromatic and aliphatic bonds about 0.05. With kappa free, N-CA is a bond like the peptide bond. Single bonds to oxygen and bonds to hydrogen are negative: at their midpoints there is less density than the superposition of independent atoms places there, as expected of a polar bond whose electrons lie towards the oxygen or the nitrogen. kappa is 1.010 and 1.013 for N, 0.983 and 0.982 for O, 0.996 and 0.995 for C in the two folds.

The definition does not place every class, and we report those it does not. The charge of the main-chain C=O bond depends on the description of the oxygen atom (+0.02 and +0.07 without kappa, -0.39 and -0.28 at the upper bound of the width with it). The side-chain C-N classes (amide, guanidinium, amine, the proline N-CD bond) disagree between the folds or reach the bound of the width; these side chains are often disordered and their bonds are few. The main-chain N-H class is +0.02 in one fold and +0.30 in the other. These are the classes on which the description of the atoms, not of the bond, is still open.

A class of the bond layer is therefore a chemical environment, fixed before the fit. With every class in its place the bond layer alone is still behind the multipole in 565 of 650 proteins.

### 4.4 Atom-centred residual density

The density that the bond layer does not reach is centred on the atoms. This was established by reading bonds out of the difference density of the IAM on the models as deposited, with one class per pair of elements; it is summarised here, and Supplementary sections SI-2 to SI-6 give it in full.

Midpoints of bonded pairs were scored against midpoints of non-bonded pairs of the same entry. For the main chain the bonds are there: the C-C and C-N read-outs are 0.643 and 0.618 over the tier, above 0.5 in 94 and 89 per cent of proteins. But the read-out falls with the B of the site, and it does not stop at 0.5, where a lost signal would stop: C-C goes from 0.696 at B below 4 A^2 to 0.476 at 17 to 25 A^2, and C-O from 0.618 to 0.317. A value below 0.5 is a signed effect. Its origin is seen when the difference density is plotted against the distance from the nearest atom. At mobile sites it is -1.15 standard deviations of the map at 0.55 A from the atom centre and returns to zero at 1.4 A, with no positive shell outside (figure 4a), and true bonds follow the same curve as empty points. The residual is centred on the atoms and not on the bonds.

**Figure 4.**
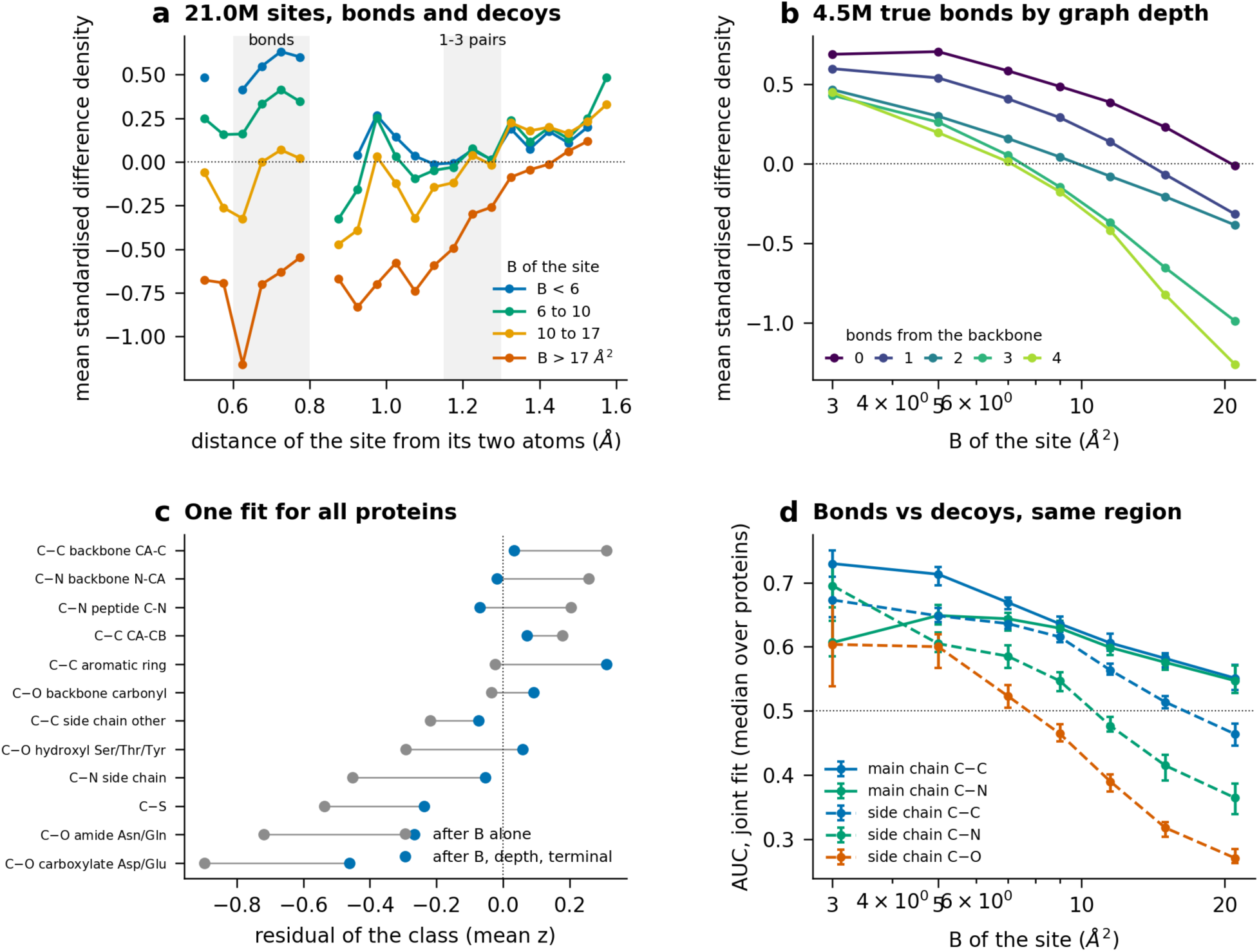
The atom-centred negative well of disordered sites. Every protein has total weight one. (a) Mean standardised difference density against the distance of the site from its two atoms, in four classes of B of the site, for all sites in the covalent range (21.0 million, bonds and decoys); the positions of bonds and of 1-3 pairs are marked. (b) True bonds (4.5 million): mean standardised difference density against B of the site (logarithmic axis), by depth of the bond in the covalent graph, 0 to 4 bonds from the backbone. (c) Residual of each chemical class (mean z) after B alone and after B, depth and the terminal term, one fit for all proteins. (d) AUC of the joint fit, bonds against decoys of the same region, against B of the site for main-chain and side-chain bonds (`fig9_negative_well`).

The term can be measured where it is centred. Writing the difference map as a sum of one radial function per atom gives a sparse linear system, which was solved in every entry (8.3 million atoms, seventeen numbers each, nothing transferred). The solution is one function that changes sign with order: at the centre of the atom +0.70 standard deviations of the map for B below 4 A^2 and -0.93 above 25 A^2 (figure 5a), with an integral of +0.10 to +0.12 electrons per ordered atom and -0.41 electrons per atom above 25 A^2. An error in the width of the atomic Gaussian would leave the integral near zero. What is found is a net excess at ordered atoms, largest on the backbone N and CA, and a net deficit at mobile ones, largest on side-chain O, N and S (figure 5e, f). With the solved functions subtracted, no bond type lies below 0.5 in any class of B: at B of 17 to 25 A^2 C-C goes from 0.421 to 0.527, C-N from 0.381 to 0.543 and C-O from 0.309 to 0.531 (figure 5b), and the bonds of residues that are not protein are read as well as those of the protein (figure 5d). The bonds had been hidden by the atoms.

**Figure 5.**
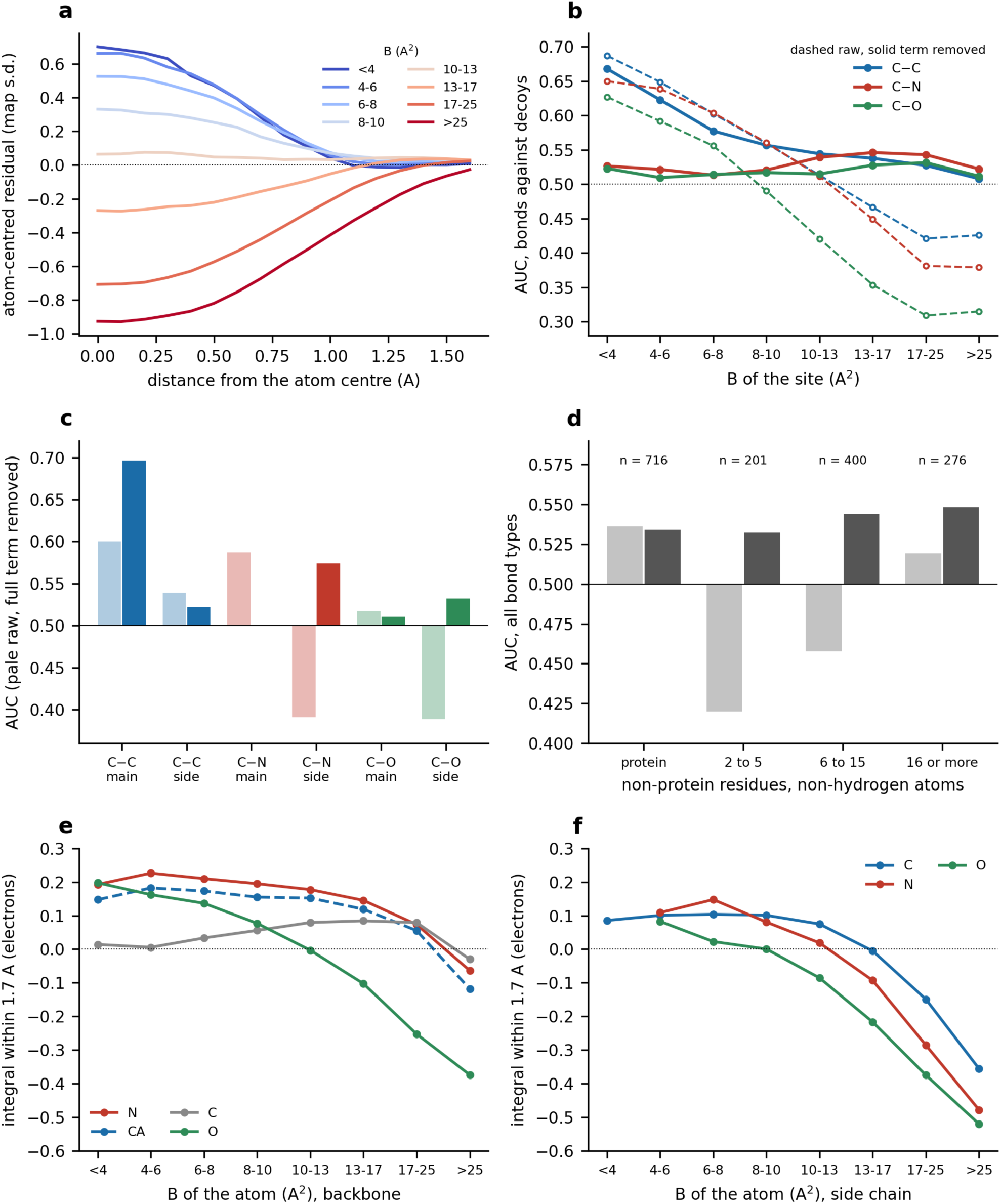
The atom-centred term, solved and removed. (a) Radial function of the difference density around protein atoms after the joint solution, in units of the standard deviation of the map of the entry, for eight classes of the atomic B (median over proteins). (b) AUC of true bonds against decoys by class of the site B, raw density (dashed) and with the atom-centred term removed (solid), for C-C, C-N and C-O. (c) The same two scores for main-chain and side-chain bonds. (d) All bond types inside non-protein residues of 2 to 5, 6 to 15 and 16 or more non-hydrogen atoms, and inside the protein. (e) Integral of the solved radial function within 1.7 A, in electrons per atom, for the backbone atoms N, CA, C and O by class of B; a point is drawn when at least fifty proteins contribute. (f) The same for side-chain C, N and O. Every protein counts once (`fig13_atom_term`).

### 4.5 Prediction of the atom-centred term from the atomic environment

The term is a property of the individual atom. Element and B do not fix it: inside an entry the logarithm of B explains a tenth of the variance of the value at the centre. At equal B the residual falls with the number of bonds between the atom and the backbone and falls again for terminal atoms (figure 4b), along the same axis on which alternative conformations, partial occupancy and missing atoms rise; packing and rigid-body motion account for more than half of the variance of B itself (Supplementary section SI-5). No correction by such named terms removes the effect, because they describe the place of a site and move bonds and empty points together (Supplementary section SI-5.4). Two properties of the instrument were learnt in passing and apply to any benchmark of this kind: the overall B must be fitted continuously to the R optimum and never tuned on the read-out (Supplementary section SI-3), and the unit of replication is the independent deposition, not the dataset (Supplementary section SI-6).

The term can be computed from the surroundings of the atom. Each atom was described by the numbers of atoms of six kinds in twenty distance shells out to 8 A, with no density and no B, and a linear model of the solved functions was fitted on half of the proteins. In the other half, subtracting the predicted functions takes the main-chain C-C read-out from 0.601 to 0.639, where the solved functions give 0.696, and the side-chain C-N read-out from 0.391 to 0.527, where they give 0.574 (Supplementary section SI-4.5). A linear count of neighbours thus recovers between two fifths and three quarters of what the full solution recovers, in proteins that took no part in the fit.

The density that the bond layer does not reach is thus centred on the atoms, belongs to the individual atom and can be computed from the positions and kinds of the atoms around it before any density is measured. These are the properties of the neighbourhood field (section 2.4); the surroundings of an atom are also what changes when a ligand binds.

### 4.6 Comparison with the transferable multipole model

With the neighbourhood field the model is better than the transferable multipole model in four proteins out of five, with fewer parameters (table 7).

**Table 7.** CEL with the neighbourhood field, hydrogens placed (650 proteins, 1,996 entries, none failed; lambda chosen on the other half of the proteins). The bond layer is the one with classes by chemical environment and kappa (159 parameters); the last three rows are the same fields on the bond layer by element pair (104 parameters).

| **model** | **parameters of the field** | **against IAM** | **against CEL** | **against multipole** | **against multipole + CEL layer** |
|---|---|---|---|---|---|
| CEL + field, matrix, one channel | 3,055 | 595 : 55, -0.0029 | 582 : 68, -0.0021 | 545 : 105, -0.0012 | 508 : 142, -0.0009 |
| CEL + field, matrix, three channels and log B | 9,221 | 601 : 49, -0.0033 | 591 : 59, -0.0024 | 557 : 93, -0.0015 | 540 : 110, -0.0012 |
| CEL + field, neural network (reference) | 96,658 | 621 : 29, -0.0033 | 617 : 33, -0.0025 | 590 : 60, -0.0016 | 581 : 69, -0.0013 |
| bond layer by element pair: one channel | 3,055 | 595 : 55, -0.0027 | 589 : 61, -0.0022 | 534 : 116, -0.0010 | 488 : 162, -0.0008 |
| bond layer by element pair: three channels and log B | 9,221 | 597 : 53, -0.0032 | 594 : 56, -0.0027 | 548 : 102, -0.0014 | 530 : 120, -0.0012 |
| bond layer by element pair: neural network | 96,658 | 621 : 29, -0.0033 | 619 : 31, -0.0027 | 582 : 68, -0.0015 | 567 : 83, -0.0012 |

The field lowers R three times more than the bond layer on which it sits (figure 6a), and the two together are better than the transferable multipole in 545 of 650 proteins with 3,214 parameters against 4,768 (figure 6b). Tripling the parameters of the matrix form moves the count from 545 to 557; a neural network with thirty times more moves it to 590. The matrix form, which is the explicit one, is what we propose.

**Figure 6.**
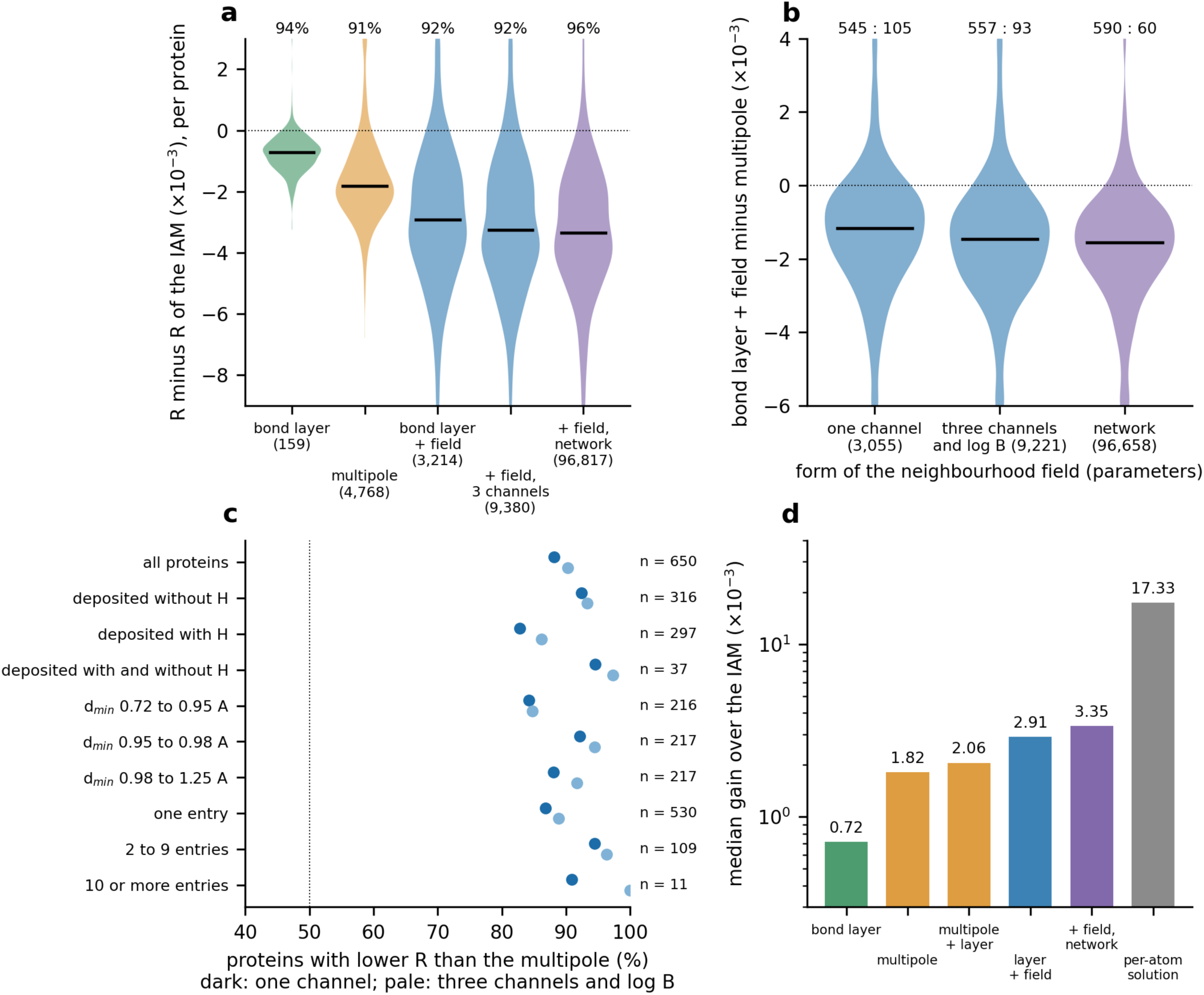
Forward benchmark with hydrogens placed: 650 proteins never seen in training, each once. (a) R minus R of the IAM (x 10^-3; below zero is better) for the bond layer, the multipole, and the bond layer with the three forms of the neighbourhood field; number of parameters in brackets; black bar: median; over each distribution the share of proteins with lower R; clipped at -9 and +3. (b) Bond layer + field minus multipole for the three forms of the field; over each distribution, better : worse; clipped at -6 and +4. (c) Share of proteins in which bond layer + field has the lower R than the multipole, lambda fixed at 1.5: all proteins, and by deposited hydrogens, thirds of resolution and number of entries of the protein; dark: one channel, pale: three channels and log B; n: proteins. (d) Median gain over the IAM (logarithmic axis); the per-atom solution is fitted on each entry and read on the reflections of the fit (section 4.8) (`fig15_forward`).

The separation of the bond classes carries through to the complete model. On the bond layer by element pair the three fields stood at 534, 548 and 582 against the multipole. Protein by protein, the complete model with one channel is better on the bond layer by chemical environment than on the layer by element pair in 480 of 650 (median -0.0003; figure 3c); with three channels in 448, and with the network in 432, by 0.0001: a field with more parameters makes up part of what the cruder bond layer leaves.

The gain of the field is not the hydrogens: on the bond layer by element pair, for which both conditions were computed, it is the same against the IAM on the models as deposited (-0.0030) and with hydrogens placed (-0.0027), while the multipole fell from -0.0043 to -0.0018.

### 4.7 Controls: training data, overlap with the multipole and counting of proteins

Three controls test the comparison of table 7: the amount of training data, the overlap of the field with the multipole, and the way proteins are counted.

#### Training data

The multipole is trained on T1 (102 usable entries). The field of table 7 is trained on one half of the T2 proteins and evaluated on the other half, about ten times more entries. We therefore trained the field on T1 alone, the training set of the multipole, with the number of updates of the main run (115 epochs), and evaluated it on all of T2 (1,996 entries, 650 proteins, none failed). On the bond layer by chemical environment the field so trained is ahead of the multipole in 414 of 650 proteins (median dR -0.0003) with 3,055 parameters, in 476 (-0.0008) with 9,221 parameters, and in 420 (-0.0004) as a network; against the IAM the three give 610 : 40, 584 : 66 and 591 : 59. On the bond layer by element pair the same control gave 403, 488 and 408. With equal training data the model is ahead of the multipole, by a smaller margin than in table 7 (545 : 105, -0.0012). The form of the model accounts for about a third of its lead; the rest comes from the larger training set, which a model with few parameters can use and a model fitted per atom type with thousands of parameters is not usually given. Both statements describe the model: it was built with few parameters so that the whole archive can train it. The number of updates matters as much as the number of entries. With 20 epochs on T1, a sixth of the updates, the field is behind the multipole with one channel (218 : 432, +0.0005) and as a network (278 : 372, +0.0002), and ahead only with 9,221 parameters (380 : 270, -0.0003); on the bond layer by element pair the three counts were 211, 248 and 361.

#### Overlap with the multipole

On the models as deposited, a field trained on the residual of the multipole with the CEL layer lowers R further in 586 of 650 proteins (-0.0019), eight times the gain of the CEL layer on the same base (569 : 81, -0.0002). The field therefore describes density that the multipole model does not. With hydrogens placed the result holds (647 proteins, 1,990 entries): the field lowers R below the multipole with the CEL layer in 570 proteins and raises it in 77 with one channel (-0.0018), 566 : 81 with 9,221 parameters (-0.0019) and 597 : 50 with the network (-0.0020), against 536 : 111 (-0.0002) for the CEL layer on the same base.

#### Counting of proteins

With lambda fixed at 1.5 for every protein, the count of CEL + field against the multipole is 573 : 77 by protein, 572 : 78 collapsing by batch first, and 1471 : 140 by entry (650 proteins, 1,611 entries); 292 : 24 for proteins deposited without hydrogens, 246 : 51 for those with, and 35 : 2 for those with both; 182 : 34, 200 : 17 and 191 : 26 in the three thirds of resolution (0.72 to 0.95, 0.95 to 0.98 and 0.98 to 1.25 A); 460 : 70, 103 : 6 and 10 : 1 for proteins with one, two to nine, and ten or more entries (figure 6c). The median dR lies between -0.0010 and -0.0012 in every one of these splits; the lead is smallest in the highest third of resolution. With lambda chosen on the other half of the proteins, as in table 7, the count by protein is 545 : 105. On the models as deposited, with the bond layer by element pair, the same splits disagreed with each other: the count by protein was 258 : 392, by entry 1020 : 591 because one protein with 581 entries decided it, and the two hydrogen classes gave 13 : 321 and 244 : 62. The disagreement was the hydrogens.

On the models as deposited the bond layer adds nothing measurable on top of the field (IAM + field -0.00307, CEL + field -0.00309): in R the field covers what the bond layer describes. The bond layer is kept because it is the part of the model that names interactions, and because it is the part that a binding calculation can address contact by contact.

### 4.8 Upper bound of the spherical correction

On each entry the radial function of every atom can be solved from the difference map itself (17 numbers per atom, regularised least squares). Put into the model, this solution lowers R by 0.0173 against the IAM in 650 of 650 proteins, and by 0.0157 against the multipole (figure 6d). It is fitted on the entry and is not a transferable model. This number is not a ceiling of what can be predicted, because the solution is fitted to the same reflections on which R is read. We repeated the solution on difference maps made from 90 per cent of the reflections and read R on the remaining 10 per cent (48 entries of the training tier, the IAM as the base, one lambda for all entries). On the reflections of the fit R falls by 0.0105; on the reflections held out of it R falls by 0.0022 (lower in 36 entries, higher in 12). About four fifths of the gain of a per-atom fit is therefore the fit to its own data. What is left, about 0.002, is the size of what the field predicted from distances already delivers on proteins it has not seen (table 7: 0.0021 below the bond layer). The spherical form, as it is predicted now, is close to what the data support, and the next gain has to come from a different form. Giving every atom three directional functions next to its spherical one raises the held-out gain of the per-atom solution to 0.0028 (lower in 42 of 48 entries), 0.0004 more than the spherical solution; a fit with 68 numbers per atom does not resolve a difference of that size, and whether direction carries density has to be asked of a model whose parameters are shared between atoms. This is answered in section 4.9: a shared-parameter field given a direction for every atom does lower R on unseen proteins beyond its spherical part, by about a fifth of the spherical term, while a bond-charge output it is trained to produce recovers nothing.

### 4.9 Environment dependence of bond charges

The charges of the bond layer respond to the surroundings of the bond, as a secondary effect about ten times smaller than the effect on the atoms. The field acts on functions centred on the atoms, and the bond layer under it is fixed: every bond of a class carries the same charge wherever it stands. Bonding electrons are the loosely held part of the cloud, so they are the part one would expect the surroundings to shape first. In the order of the model, atoms and bonds are built and refined first and the field is the last layer; if the surroundings act on the bonds, that action belongs to the field. Before building it we measured how much there is to predict, that is, what is left on the bonds once atoms and bonds are in place.

On the residual map of the bond layer every covalent bond was given three free quantities: a change of its charge, taken from or returned to the valence shells of its two atoms so that no electron is added; a dipole along the bond axis; and a dipole across it (four numbers per bond; the dipoles are the first order of a shifted bond charge). They were solved for all bonds of the crystal together, alone and together with the atom functions of section 4.8 with direction (68 numbers per atom), on maps made from 90 per cent of the reflections, and R was read on the other 10 per cent (109 entries of the training tier, 305,983 atoms and 282,434 bonds; base: the bond layer; one weight, 0.5, for every entry and every row).

| **freed on the residual of the bond layer** | **numbers** | **R on held-out reflections: lower : higher, median change** | **R on the reflections of the fit, median change** |
|---|---|---|---|
| charge of the bonds | 1 per bond | 83 : 26, -0.00017 | -0.0013 |
| dipoles of the bonds | 3 per bond | 49 : 60, +0.00007 | -0.0012 |
| charge and dipoles of the bonds | 4 per bond | 67 : 42, -0.00026 | -0.0026 |
| atom functions with direction | 68 per atom | 90 : 19, -0.00214 | -0.0157 |
| atom functions and bonds together | 68 per atom, 4 per bond | 92 : 17, -0.00226 | -0.0159 |
| the same against the atom functions alone |  | 85 : 24, -0.00012 | -0.0004 |

Three things follow. The bonds do respond: the charge of a bond, once freed, lowers R on reflections it was not fitted to in 83 of 109 entries, and bonds added to the atom functions lower it further in 85. The response is small: 0.0002 to 0.0003 against the 0.0022 of the atom functions, about one tenth, with a change of 0.06 electron per bond (r.m.s.), which is of the size of the class charges themselves (0.03 to 0.3 electron). And it is a response of amount, not of position: the dipoles lower R on the reflections of the fit as much as the charge does, and not at all on the reflections held out. The constraint matters. Without it the solution moves 0.28 electron per bond, the gain of the bonds alone on held-out reflections is no larger (0.0002), and most of what a free per-bond solution finds is noise; the numbers above are with a ridge that holds the change at 0.06 electron (Methods).

The effect of the surroundings on the bonding electrons is therefore real and secondary. For the atoms the field is the main term of the model. For the bonds it is a correction of the bond charge about ten times smaller, and the data support no displacement of the bond. This is a bound and not yet a model: the solution is fitted on the entry, as in section 4.8, and the field that predicts the change of bond charge from the surroundings has not been trained. One of the largest entries of the tier was not finished when this was written and one has no data; they are not in the table.

A field of shared parameters was then trained to produce these solutions: the neighbourhood field of section 4.6 with two outputs added, a direction for every atom and a change of charge for every bond. Trained on one half of the proteins and read on the other (the 650 unseen of section 4.6), it separates the two effects. Direction gives a small, consistent gain: on top of the spherical part it lowers R in 593 of 650 proteins, by a further 0.00015; this is the first directional gain on proteins the field was not trained on, about a fifth of the spherical term. The bond charge gives nothing: the field’s predicted change of bond charge correlates with the solved one at only 0.04, and adding it lowers R in 196 of 650 proteins (median +0.00002). This field, whose target is the harder joint solution, is weaker overall than the field of section 4.6 (its spherical part lowers R by 0.00073 against the bond layer, against 0.0021 in table 7) and does not beat the multipole; it is an existence proof for a directional term, not a new main result. A field with direction as strong as the spherical term, with the bond charge left to the bond layer where the data keep it, is the next model.

### 4.10 Inverse read-out of bond density

Read without R, the density shows the bonds that a compound design calculation addresses: the bonds of ligands and the bonds and contacts of the pocket around them. The polar bonds of ligands read below empty points in the density left by independent atoms, and the neighbourhood field predicted from the surroundings turns them the right way round, to the values that the per-atom solution of the entry gives. The read-out also confirms the bond classes: the classes by chemical environment take up bond density that one class per element pair cannot.

Every covalent bond midpoint and every contact site of the 2,006 entries is a site, named by the classes of section 2.3. A site is compared with empty points of the same entry that are matched to it on the distance to the nearest atom, the distance to the nearest hydrogen, ln B and occupancy. The AUC of a class in an entry is the fraction of matched points that its sites exceed: 0.5 means that a bond cannot be told from an empty point in the same place, and a value below 0.5 that it carries less density than such a point. An entry counts for a class when it has at least five matched sites, a protein by the median over its entries (650 proteins; 1,927 of the 2,006 entries gave a read-out). Three values are read at every site and its matched points alike: the difference density after the atoms; that density minus the neighbourhood field predicted from distances, multiplied by a factor lambda; and that density minus the per-atom solution of the entry. Sites are labelled by their place relative to ligands. A ligand bond lies inside a residue that is neither protein nor water and has at least six non-hydrogen atoms, a drug-like compound or a cofactor; a pocket site is any other bond or contact within 4.5 A of a heavy atom of such a ligand, symmetry images included. All sites together are the control.

The factor lambda was calibrated on these sites against the per-atom solution, which removes the atom-centred term of each entry using the entry’s own data. Over the ligand classes, the protein-weighted mean difference between the AUC after the predicted field and the AUC after the per-atom solution falls from 0.069 without the field to 0.023 at lambda = 1.5 and rises beyond it (0.037 at 2, 0.073 at 3; figure 7a). The factor of the forward benchmark (section 4.7), chosen on R, is thus the factor at which the field predicted from the surroundings reproduces the measured atom term on ligand bonds. At the pocket the difference is flat between lambda 0.75 and 1.5 (0.078 to 0.080); over all sites it is lowest at 1.0 (0.051, against 0.066 at 1.5); with the closer matching on both nearest atoms the ligand optimum is 1.0, with 1.25 next. Removing the field of the hydrogen atoms makes the agreement worse for every group of sites (at lambda 1.5: ligand 0.037, pocket 0.112, all sites 0.069): the field on hydrogens is part of what the model has to predict. A factor chosen for the largest AUC would not be a calibration: the AUC of the polar classes rises with lambda to the end of the scan while that of the C-H bond on CA falls to 0.19 at lambda = 3.

**Figure 7.**
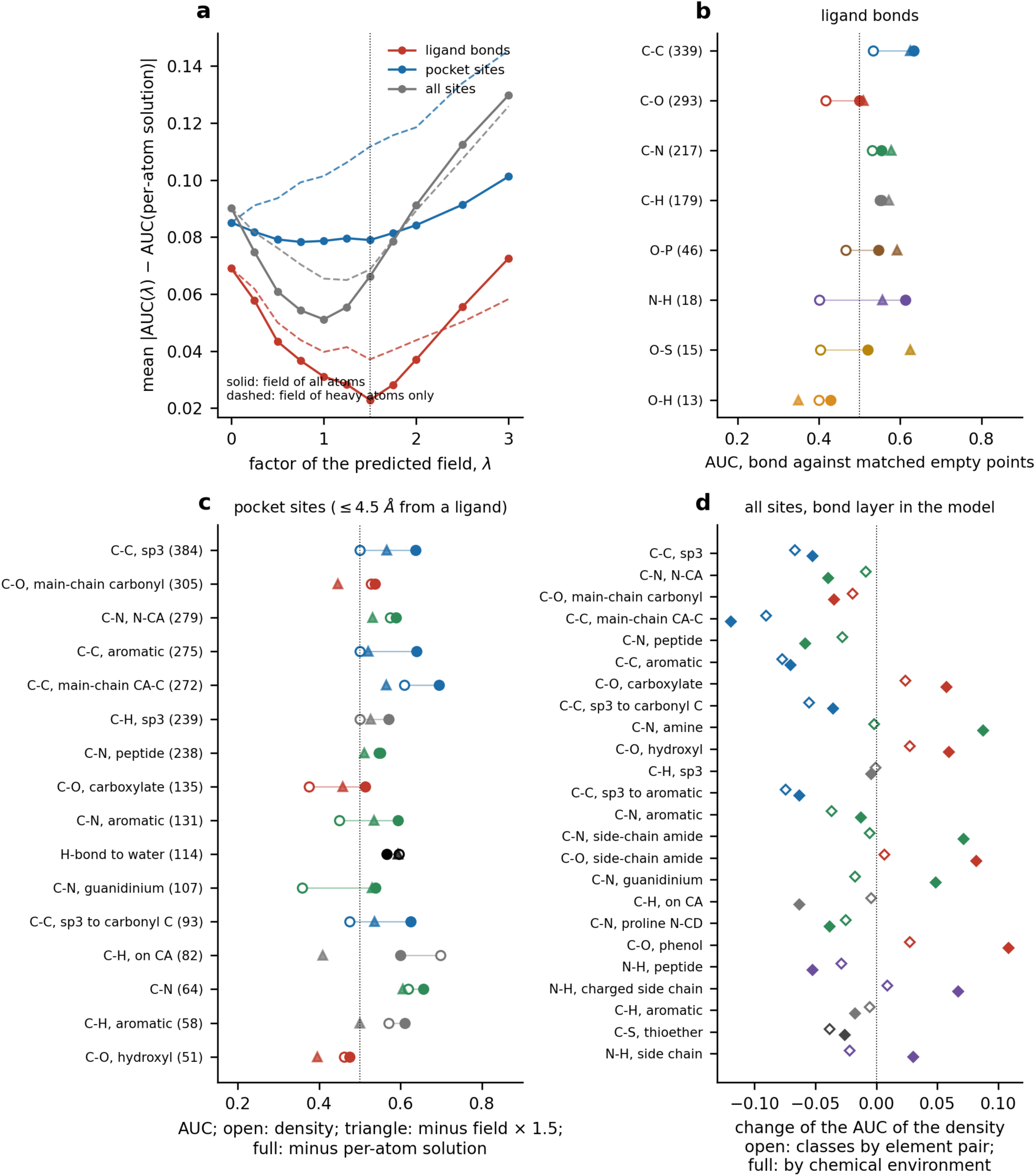
Reading bonds out of the density for compound design, hydrogens placed (section 4.10). AUC of bond midpoints and contact sites against matched empty points: median over proteins of the per-protein median, map after the atoms. (a) Calibration of the factor lambda of the predicted neighbourhood field: protein-weighted mean absolute difference between the AUC after subtracting the field times lambda and the AUC after subtracting the per-atom solution of the entry, for ligand bonds, pocket sites (within 4.5 A of a ligand) and all sites; solid: field of all atoms, dashed: field of heavy atoms only; dotted line: lambda = 1.5. (b) Ligand bonds (classes with at least 10 proteins) and (c) pocket sites (at least 50 proteins): density (open circle), density minus the predicted field at lambda = 1.5 (triangle) and density minus the per-atom solution (full circle); number of proteins in brackets; dotted line: 0.5. (d) All sites (at least 100 proteins): change of the AUC of the density when the bond layer is in the model, with classes by element pair (open) and by chemical environment (full) (`fig17_inverse_h`).

Ligand bonds (figure 7b). The C-C bonds of ligands lie above their matched points in 200 proteins and below in 135 in the density, in 257 and 78 with the predicted field removed, and in 298 and 36 with the per-atom solution removed. The polar bonds of ligands read the wrong way round in the density: C-O at 0.417 (75 : 216), the phosphate O-P bond at 0.466 and the sulfonyl and sulfate O-S bond at 0.404. With the predicted field removed they reach 0.510, 0.593 and 0.625, at or above the per-atom solution (0.500, 0.546 and 0.521). These are the bonds through which a compound makes its polar contacts. In the density left by independent atoms they carry less density than an empty point at the same distance from an atom, and the field computed from the positions and kinds of the surrounding atoms alone restores them. The hydroxyl O-H bond of ligands stays below 0.5 under all three values (13 proteins).

Pocket sites (figure 7c). Around the ligand the carboxylate C-O and guanidinium C-N bonds, the groups that form salt bridges with ligands, read the wrong way round in the density (0.375 and 0.358) and are restored by the predicted field (0.458 and 0.530) and by the per-atom solution (0.513 and 0.539); the aromatic C-N bond goes from 0.450 to 0.535. Hydrogen bonds to water in the pocket are read at 0.597 and are unchanged by the field (0.594). Two pocket classes are taken too far by the field at lambda = 1.5: the C-H bond on CA (0.698 in the density, 0.408 with the field, 0.600 with the per-atom solution) and the main-chain carbonyl (0.528, 0.446 and 0.537). Both are main-chain bonds; the field predicted around the main chain is the part of the model to refine for the pocket.

Over all sites the same pattern holds (table 8, lower part). The bonds of the main chain are read directly: CA-C, N-CA and the peptide bond lie above their matched points in 568, 528 and 499 proteins, and the C-H bond on CA and the main-chain N-H bond are the most visible of all (0.73 and 0.72). The bonds from side chains to nitrogen and the carboxylate C-O bond are read the wrong way round: the amine C-N bond lies below its matched points in 501 proteins and above in 20. These are not bonds without density. When the predicted field is subtracted from site and matched point alike the amine bond is above in 501 proteins, the side-chain amide and guanidinium C-N bonds in 444 and 401, and the sp3 C-C bond rises from 370 to 544; with the per-atom solution every carbon-carbon class is above 0.5 in more than four proteins of five (CA-C 609 : 18). The read-out below 0.5 is therefore the atom-centred term of section 4.4 seen from a bond midpoint, and its sign depends on the class of the bond. At lambda = 1.5 the predicted field takes the C-H bond on CA and the main-chain N-H bond below 0.5 (0.400 and 0.389); with the field as predicted (lambda = 1) both read 0.500, and the per-atom solution gives 0.610. The factor that suits the ligand bonds removes too much around the main chain.

**Table 8.** Reading bonds out of the density, hydrogens placed (median over proteins of the per-protein median AUC; in brackets proteins above : below 0.5; field subtracted at lambda = 1.5). Upper part: bonds of ligands (classes with at least 10 proteins) and sites of the pocket within 4.5 A of a ligand (classes with at least 50 proteins). Lower part: all sites (classes with at least 100 proteins), with the AUC of the density when the bond layer is in the model. Every class at every factor of the scan is in the source data of figure 7.

| site | class | proteins | density | minus field (x 1.5) | minus per-atom solution |  |
|---|---|---|---|---|---|---|
| ligand | C-C | 339 | 0.534 (200 : 135) | 0.625 (257 : 78) | 0.633 (298 : 36) |  |
| ligand | C-O | 293 | 0.417 (75 : 216) | 0.510 (158 : 130) | 0.500 (145 : 142) |  |
| ligand | C-N | 217 | 0.531 (134 : 80) | 0.578 (155 : 60) | 0.554 (153 : 61) |  |
| ligand | C-H | 179 | 0.554 (97 : 78) | 0.571 (98 : 80) | 0.550 (99 : 73) |  |
| ligand | O-P | 46 | 0.466 (18 : 26) | 0.593 (32 : 14) | 0.546 (29 : 15) |  |
| ligand | N-H | 18 | 0.401 (5 : 12) | 0.556 (10 : 8) | 0.613 (10 : 7) |  |
| ligand | O-S | 15 | 0.404 (5 : 10) | 0.625 (11 : 4) | 0.521 (8 : 7) |  |
| ligand | O-H | 13 | 0.400 (4 : 8) | 0.350 (3 : 10) | 0.429 (5 : 8) |  |
| pocket | C-C, sp3 | 384 | 0.500 (189 : 188) | 0.566 (273 : 107) | 0.637 (340 : 40) |  |
| pocket | C-O, main-chain carbonyl | 305 | 0.528 (170 : 129) | 0.446 (108 : 187) | 0.537 (183 : 114) |  |
| pocket | C-N, N-CA | 279 | 0.575 (208 : 67) | 0.531 (164 : 105) | 0.589 (206 : 64) |  |
| pocket | C-C, aromatic | 275 | 0.500 (134 : 134) | 0.521 (157 : 112) | 0.639 (241 : 31) |  |
| pocket | C-C, main-chain CA-C | 272 | 0.609 (214 : 50) | 0.565 (181 : 86) | 0.695 (247 : 22) |  |
| pocket | C-H, sp3 | 239 | 0.500 (110 : 118) | 0.526 (124 : 110) | 0.571 (129 : 105) |  |
| pocket | C-N, peptide | 238 | 0.550 (151 : 85) | 0.510 (125 : 107) | 0.547 (160 : 68) |  |
| pocket | C-O, carboxylate | 135 | 0.375 (24 : 111) | 0.458 (50 : 82) | 0.513 (70 : 56) |  |
| pocket | C-N, aromatic | 131 | 0.450 (46 : 79) | 0.535 (78 : 50) | 0.594 (99 : 29) |  |
| pocket | H-bond to water | 114 | 0.597 (77 : 37) | 0.594 (76 : 38) | 0.566 (82 : 29) |  |
| pocket | C-N, guanidinium | 107 | 0.358 (17 : 87) | 0.530 (59 : 44) | 0.539 (67 : 34) |  |
| pocket | C-C, sp3 to carbonyl C | 93 | 0.475 (38 : 53) | 0.536 (52 : 38) | 0.625 (75 : 16) |  |
| pocket | C-H, on CA | 82 | 0.698 (55 : 23) | 0.408 (32 : 48) | 0.600 (46 : 35) |  |
| pocket | C-N | 64 | 0.620 (49 : 15) | 0.606 (45 : 19) | 0.656 (55 : 9) |  |
| pocket | C-H, aromatic | 58 | 0.571 (33 : 22) | 0.500 (27 : 28) | 0.611 (37 : 18) |  |
| pocket | C-O, hydroxyl | 51 | 0.462 (18 : 31) | 0.396 (12 : 38) | 0.475 (18 : 31) |  |
| **class (all sites)** | **proteins** | **density** | **minus field (x 1.5)** | **minus per-atom solution** | **density, layer by element pair** | **density, layer by chemical environment** |
| C-C, sp3 | 639 | 0.513 (370 : 267) | 0.565 (544 : 95) | 0.626 (609 : 30) | 0.446 | 0.460 |
| C-N, N-CA | 629 | 0.592 (528 : 99) | 0.532 (400 : 222) | 0.577 (512 : 115) | 0.583 | 0.552 |
| C-O, main-chain carbonyl | 629 | 0.550 (424 : 203) | 0.468 (212 : 415) | 0.537 (409 : 217) | 0.531 | 0.515 |
| C-C, main-chain CA-C | 628 | 0.622 (568 : 60) | 0.572 (504 : 122) | 0.680 (609 : 18) | 0.531 | 0.503 |
| C-N, peptide | 597 | 0.577 (499 : 98) | 0.525 (376 : 219) | 0.559 (487 : 110) | 0.549 | 0.519 |
| C-C, aromatic | 592 | 0.534 (391 : 199) | 0.541 (388 : 201) | 0.623 (529 : 59) | 0.457 | 0.464 |
| **class (all sites)** | **proteins** | **density** | **minus field (x 1.5)** | **minus per-atom solution** | **density, layer by element pair** | **density, layer by chemical environment** |
| C-O, carboxylate | 564 | 0.342 (41 : 521) | 0.535 (354 : 206) | 0.499 (276 : 285) | 0.366 | 0.400 |
| C-C, sp3 to carbonyl C | 555 | 0.445 (144 : 407) | 0.537 (377 : 174) | 0.606 (479 : 73) | 0.390 | 0.410 |
| C-N, amine | 523 | 0.298 (20 : 501) | 0.717 (501 : 17) | 0.550 (363 : 156) | 0.296 | 0.385 |
| C-O, hydroxyl | 523 | 0.464 (168 : 344) | 0.450 (170 : 347) | 0.503 (267 : 248) | 0.491 | 0.523 |
| C-H, sp3 | 516 | 0.508 (265 : 245) | 0.538 (279 : 231) | 0.551 (296 : 213) | 0.507 | 0.503 |
| C-C, sp3 to aromatic | 512 | 0.560 (375 : 130) | 0.469 (185 : 320) | 0.621 (433 : 71) | 0.485 | 0.497 |
| C-N, aromatic | 509 | 0.462 (191 : 311) | 0.584 (386 : 117) | 0.589 (411 : 88) | 0.425 | 0.450 |
| C-N, side-chain amide | 508 | 0.383 (66 : 439) | 0.630 (444 : 63) | 0.559 (372 : 129) | 0.377 | 0.454 |
| C-O, side-chain amide | 501 | 0.375 (45 : 446) | 0.527 (308 : 188) | 0.507 (263 : 228) | 0.382 | 0.457 |
| C-N, guanidinium | 495 | 0.379 (73 : 419) | 0.600 (401 : 90) | 0.563 (381 : 111) | 0.361 | 0.427 |
| C-H, on CA | 412 | 0.730 (320 : 83) | 0.400 (137 : 267) | 0.610 (264 : 142) | 0.726 | 0.667 |
| C-N, proline N-CD | 376 | 0.543 (238 : 123) | 0.600 (291 : 76) | 0.583 (279 : 90) | 0.518 | 0.505 |
| C-O, phenol | 326 | 0.410 (78 : 239) | 0.492 (144 : 169) | 0.417 (67 : 251) | 0.438 | 0.519 |
| N-H, peptide | 286 | 0.719 (212 : 66) | 0.389 (103 : 178) | 0.610 (171 : 108) | 0.690 | 0.667 |
| N-H, charged side chain | 280 | 0.433 (114 : 154) | 0.667 (187 : 81) | 0.500 (131 : 132) | 0.442 | 0.500 |
| C-H, aromatic | 239 | 0.517 (125 : 105) | 0.455 (108 : 125) | 0.571 (128 : 104) | 0.512 | 0.500 |
| C-S, thioether | 216 | 0.411 (53 : 160) | 0.565 (143 : 65) | 0.579 (158 : 51) | 0.373 | 0.385 |
| N-H, side chain | 181 | 0.500 (83 : 87) | 0.625 (119 : 58) | 0.579 (104 : 71) | 0.478 | 0.530 |

With the bond layer in the model the read-out shows what the layer has taken up (figure 7d). Under the layer by element pair the carbon-carbon classes fall towards 0.5 (CA-C from 0.622 to 0.531) and the nitrogen and oxygen classes read the wrong way round do not move (amine 0.298 to 0.296, side-chain amide C-N 0.383 to 0.377): one number for all C-N bonds cannot serve the peptide bond and the amine. Under the classes by chemical environment these classes move towards 0.5 (amine 0.385, amide 0.454, guanidinium 0.427) and the main-chain classes are taken up further (peptide bond 0.519, N-CA 0.552, CA-C 0.503). The inverse read-out uses no R and no training label, and it gives the fourth result again: bonds that one class had put together are different in the density. The field subtracted in table 8 is the one-channel field trained on the residual of the bond layer by element pair.

The non-covalent classes of the layer were examined on the models as deposited (Supplementary sections SI-1 and SI-4.4). Against a placebo class at empty points matched on position, the carbonyl n-to-pi* class and the hydrogen-bond classes lower R more than their placebo (521 : 45 proteins for the carbonyl class), the salt bridge does not differ from it, and the C-H-pi and stacking classes are worse than their own control. What the layer reads at polar contacts is density beyond position, without specificity to the named geometry, and a contact class should be called supported by density only when it passes both the forward control and the inverse read-out.

## 5. Discussion

### 5.1 A transferable, explicit model of electron density

The task was a transferable, explicit model of electron density beyond independent atoms that a design calculation can use. Four things are now in place. A bond layer in which every number is a named interaction and electrons are conserved. A definition of its classes by chemical environment, fixed beforehand, under which the charges can be read bond by bond: the carbon skeleton positive and narrow, polar single bonds negative, and the negative C-N of the cruder definition traced to an effect on the atoms N and CA. A neighbourhood field, also explicit, that makes an atom’s cloud depend on all atoms around it; this is the part that matters in a binding site, where the surroundings of protein atoms change when a ligand arrives. And a benchmark condition, hydrogens placed by one rule, under which different models can be compared without the comparison measuring something else. Under that condition the model is better than a transferable multipole model in four proteins of five, with fewer parameters.

### 5.2 Open parts of the model

Three parts of the model remain open. The field is spherical: it describes contraction and expansion of an atom’s cloud, not its polarisation in a direction, which is what a multipole describes and what remains better in the multipole at the highest resolution. And the bond layer, although it is the interpretable part, adds little in R once the field is present; with the classes defined by chemical environment most charges read as chemistry, and the classes that do not (the main-chain carbonyl, the side-chain C-N bonds, the main-chain N-H) are those on which the description of the atom is still open. And the field does not yet act on the bonds: their charges are the same in every surrounding, although a change of about 0.06 electron per bond is supported by held-out reflections (section 4.9). The effect is a tenth of that on the atoms, so it changes the reading of individual bonds more than it changes R.

### 5.3 Limitations

The comparison with the multipole is with our own implementation and our own training, not with a published library. The field of the main table was trained on more entries than the multipole (section 4.7). The rule for hydrogens places rotatable hydrogens (hydroxyl, amine, methyl) at library positions and cannot refine them; all models share this error. The investigation of sections 4.4 and 4.5 and of Supplementary sections SI-1 to SI-6 was made on the models as deposited, with the bond layer by element pair, and has not been repeated with hydrogens placed; the read-out of section 4.10 has. lambda, the weight of the field, is one number chosen on half of the proteins; it corrects for the fact that a difference map shows a missing feature below its true height. The factor at which the predicted field best reproduces the per-atom solution on ligand bonds (1.5) takes the main-chain C-H, N-H and carbonyl bonds too far; one factor does not yet suit every class of site.

### 5.4 Outlook

For the model: a field with direction as strong as the spherical term (a first, small directional gain on unseen proteins is shown in section 4.9, while the bond charge is better left to the bond layer, where the data keep it); training the field directly on R instead of on per-entry solutions; equal weighting of proteins in training. For the use it was built for: the same two layers apply to a ligand in a pocket without change, since nothing in them is specific to proteins, and the non-protein residues of the test set are described like protein (Supplementary section SI-4.3). The next step is to compute, for complexes with known affinity, how the neighbourhood field of the pocket atoms changes on binding.

## Methods

### Hydrogens

Deposited hydrogen and deuterium atoms were removed and all hydrogens placed with gemmi 0.7.5 (prepare_topology, HydrogenChange.ReAddButWater) from the CCP4 monomer library: electron-cloud X-H distances, no hydrogens on water, occupancy and alternative-conformer label of the parent, isotropic B equal to 1.2 times the equivalent B of the parent heavy atom. Monomers without a library entry received no hydrogens and are listed in the supplementary material. The same rule was applied to the training and the test set, and every model was trained again on the training set with hydrogens.

### Conserving bond layer

Donor atoms: the two atoms of a covalent bond, half of the charge each; for a contact site, its two partners, half each, a partner that is an aromatic ring dividing its half equally among the ring atoms; a site with one partner takes all from it. The removed charge has the normalised valence form factor of the donor element and the displacement of the donor atom. Parameters (one charge and one width per class) were trained on the training set by minimising the mean R with the solvent and overall B of each entry held at their IAM values, and evaluated with scale, solvent and overall B fitted on the entry. With the classes by chemical environment, the class of a bond in a standard residue is assigned from the names of its two atoms (the rules are in the released code); one kappa per element (C, H, N, O and S, bounded between 0.85 and 1.20) scales the radial coordinate of the valence shell at fixed population, and all classes and the five kappa are fitted together.

### Neighbourhood field

Per-bond solution (section 4.9): on the residual of the bond layer, sampled at the same points as the per-atom solution, every covalent bond carries a change of charge dq (a Gaussian at the bond midpoint with the width of its class and the thermal width of its two atoms, minus half of dq taken from the unit valence density of each atom with its B) and a dipole vector p (the first derivative of the same Gaussian), resolved afterwards along and across the bond axis; symmetry images share the numbers of their parent, the vector rotated with the operator. Unknowns in units of 0.1 electron and 0.01 electron Angstrom; ridge on the bond numbers 1 (strengths 0.064, 1, 3, 10, 30 and 100 were all solved; at 0.064 the r.m.s. change is 0.28 electron, at 1 it is 0.06, at 100 it is 0.008); atom functions with direction: one spherical and three directional radial functions per atom on the 17 nodes, ridge 0.064; conjugate gradients, all atoms and bonds of the crystal together. Maps from 90 per cent of the reflections, chosen at random per entry; R on the other 10 per cent with scale, solvent and overall B fitted on them; the weight of the term on the grid 0.1, 0.25, 0.5, 0.75, 1, 1.5, 2, 3, the value 0.5 being the one at which the atom functions do best on held-out reflections, used for all rows.

Per-atom solution: for every atom a radial function on 17 nodes (0 to 1.6 A in steps of 0.1 A, zero at 1.7 A) was solved by conjugate gradients from the difference map of the entry, sampled on 64 directions per shell, with a ridge of 0.001. Field: pairs of atoms within 8 A, including symmetry mates, expanded on hat functions of the logarithm of the distance (30 nodes from 0.5 A in steps of 10%); 14 atom classes; one or three channels; six passes of the propagation; trained with Adam (rate 0.003, 20 epochs, entries in random order) to reproduce the per-atom solutions, on one half of the proteins, and evaluated on the other half. The field enters the structure factor with one weight lambda (grid 0.5, 1, 1.5, 2, 3, 4, 5) chosen on the half of the proteins not being scored. Scale, bulk solvent and overall B are fitted on the entry for every model and every lambda.

#### Data

All X-ray entries of the Protein Data Bank with deposited structure factors and d_min <= 1.00 A at the date of download (2,006; list in the supplement). Entries are read with every alternative conformer and its occupancy, anisotropic displacement parameters where deposited, and deuterium as hydrogen. Removing alternative conformers, as in an earlier reader, doubled the IAM R of 2VB1 (0.17 against 0.085) and created an apparent gain of the bond layer that was an artefact. Ten entries were not computed (unsupported element or unreadable file). Proteins are grouped at 95% sequence identity (726 groups); depositing batches are defined from the deposition records.

#### Structure factors

IAM scattering factors are the nine-parameter IT92 fits (International Tables for Crystallography, Vol. C; Prince, 2004). Structure factors are computed by direct summation over atoms and symmetry operators on GPU, in matrix form. A flat bulk-solvent term, one scale and one overall isotropic B are fitted on each entry; the overall B is refined continuously to 0.001 A^2 at the R optimum (section 3.1).

#### Training

The eight CEL-lean numbers and the ten kappa numbers were fitted on T1 in two folds by minimising the mean R of the training fold (Adam; per-entry solvent and B frozen), and evaluated on the other fold and on all T2 entries outside T1 with only scale, solvent and overall B fitted. The held-out gain is flat from 16 training structures upward. The kappa model is the spherical subset of the kappa formalism with single-zeta Slater valence functions, implemented by us. The multipole reference is a Hansen-Coppens model of our own implementation (Hansen & Coppens, 1978), explicit and transferable in exactly the same sense as CEL: for an atom of type t and element e, with *s* = sin *θ/λ* and *q* = 4*πs*,

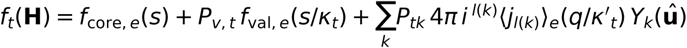

with valence population *P_v,t_*, contraction parameters *k_t_*, *k׳_t_* and multipole populations *P_tk_* for the fifteen real spherical harmonics up to *l* = 3, about 4,700 numbers over all atom types, trained once per fold and transferred with nothing fitted per entry but scale, solvent and overall B. Both CEL and the multipole model are therefore explicit, analytic and transferable; they differ in size (about ninety numbers against about 4,700) and in what each number means (a charge and width on a named bond or contact, against a multipole population on an atom type).

#### IAS-type baseline

The same Gaussian basis as CEL, one charge per bond, refined on the work reflections of the entry; evaluated on the test reflections.

#### Inverse scores

Sites: midpoints of all pairs of non-hydrogen atoms closer than the sum of covalent radii plus 1.5 A. Raw: the difference density (*F*_obs_ - *F*_IAM_, model phases) at the site. Single: the amplitude of one Gaussian of the CEL width and the thermal width of the two atoms fitted at the site alone. Joint: the amplitudes of all sites of the entry refined together. z: the raw score standardised by the median and the median absolute deviation of the covalent-range decoys of the entry. AUC: true bonds against decoys of the same entry; inside a B bin only when the entry has at least five true and five decoy sites in the bin.

#### Graph terms

depth: number of bonds from the outer atom of a bond to the nearest of N, CA, C, capped at 5. term: 1 if the bond involves an atom with one heavy-atom neighbour.

#### Statistics

Every protein counts once (median over its entries, or entry weights 1/n). Sign tests are two-sided binomial tests on proteins. For the class-ablation scan of section SI-1, which tests about twenty classes, significance is judged against a Benjamini-Hochberg false-discovery-rate threshold; the surviving classes pass at any conventional correction. Variance shares are from linear models with the named terms; missing values are set to the median.

#### Compute

Texas Advanced Computing Center, Horizon; one entry takes seconds to minutes on one GPU.

## Data availability

All structures and structure factors are public entries of the Protein Data Bank; the list of entries, the protein groups and the per-entry results are given in the supplement.

## Code availability

The code used in this work is available from the corresponding authors on reasonable request for academic, non-commercial use, under a licence that does not permit redistribution.

## Acknowledgements

Artificial-intelligence tools (a large language model) assisted with code development, data analysis and manuscript drafting; all computations were run on the authors’ resources and every result was verified by the authors. AI tools are not authors.

## Author contributions

(CRediT). Y.S.: Conceptualization; Methodology; Software; Investigation; Project administration; Supervision; Writing - original draft; Writing - review and editing. H.C.: Investigation; Resources; Writing - original draft; Writing - review and editing. H.S.: Funding acquisition; Writing - original draft; Writing - review and editing. Computing resources were provided by H.C. and the Texas Advanced Computing Center. Y.S. initiated and led the project. Y.S. is the corresponding author for TriaX AI Inc.; H.C. is the corresponding author for the Texas Advanced Computing Center. All authors read and approved the manuscript.

