## Supplementary Material for "Bonds, a neighbourhood field and missing hydrogens: a transferable electron-density model of proteins"

### Supplementary figures

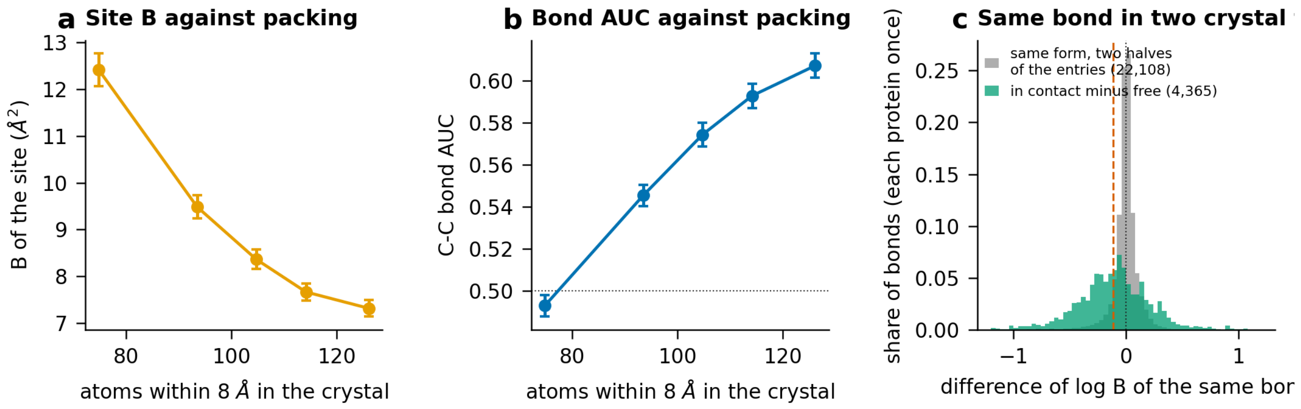


Figure S1. Packing and lattice contact as explicit terms of the site B. (a) B of the site and (b) C-C bond AUC against the number of atoms within 8 A in the crystal: the sites of each entry in five classes of packing density, mean over proteins with 95% confidence interval. (c) Difference of log B of the same bond between two crystal forms of one protein, copy in a lattice contact minus free copy (green, 4,365 bonds), against the difference between two halves of the entries of one crystal form (grey, 22,108 bonds); every protein has total weight one; dashed red line: weighted mean (`fig7_packing_contact`).

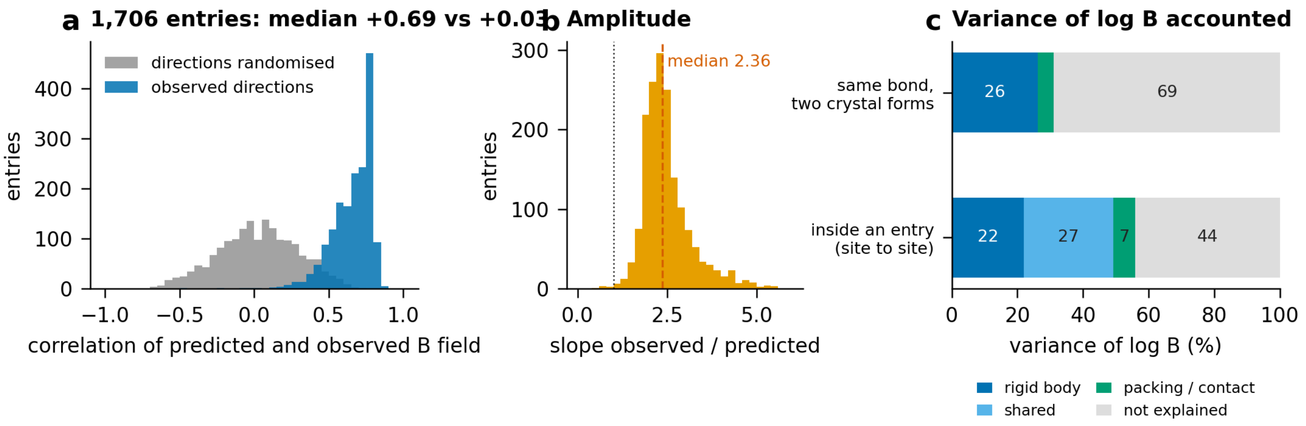


Figure S2. Rigid-body motion from the anisotropic tensors against the observed B. (a) Correlation, per entry, of the B field predicted by a rigid-body fit per chain with the observed B (1,706 entries; median +0.69), and the same with the directions randomised (median +0.03). (b) Slope of the observed on the predicted amplitude per entry; dashed red: median, 2.36; dotted: 1. (c) Variance of log B accounted for by rigid-body motion, by packing and contact, by both (shared) and by neither: among the sites of one entry, and for the same bond in two crystal forms (`fig8_rigid_body`).

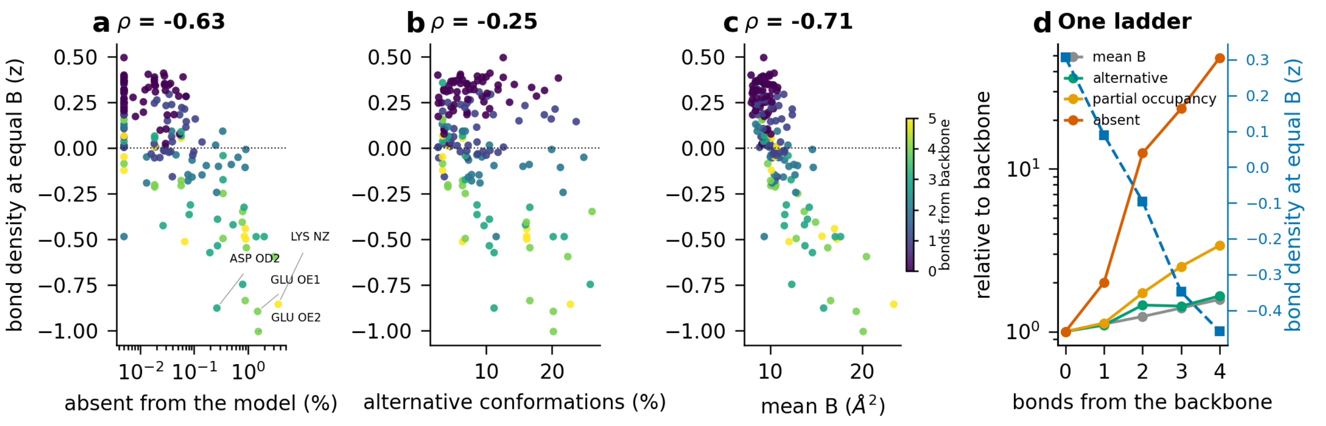


Figure S3. A ladder of disorder along the covalent graph. One point per atom position of the twenty amino acids (165 positions), coloured by the number of bonds from the backbone. Bond density at equal B (z) against (a) the fraction of atoms absent from the model (logarithmic axis), (b) the fraction modelled with alternative conformations and (c) the mean B; rho is the rank correlation, and the four positions of lowest z are named in (a). (d) Mean B, alternative conformations, partial occupancy and absence, each relative to the backbone (logarithmic axis), and the bond density at equal B (right axis), against the number of bonds from the backbone (`fig10_dynamics_ladder`).

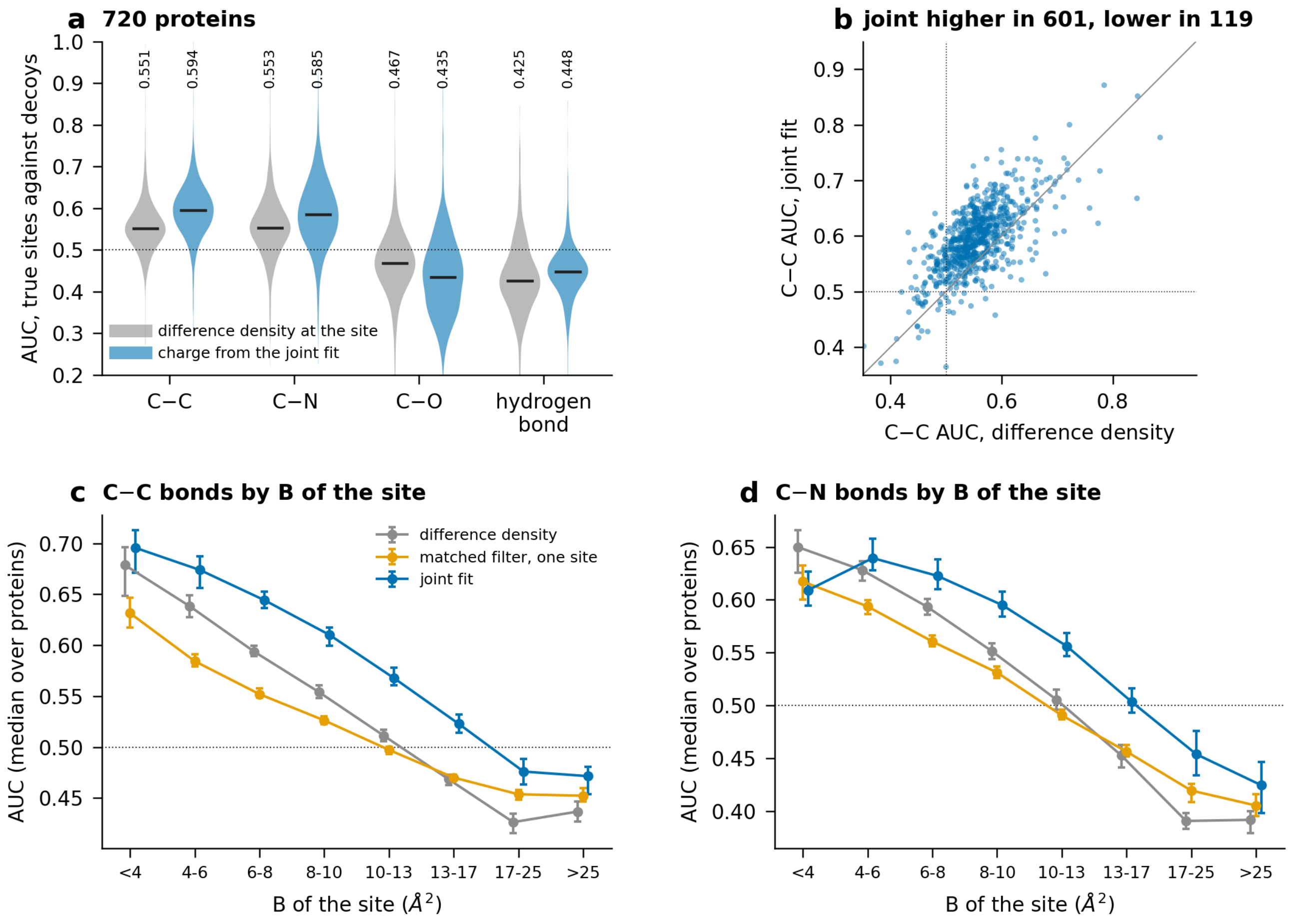


Figure S4. Inverse benchmark on the models as deposited (section SI-2). AUC of true sites against decoys, each protein once (median over its entries). (a) Distribution over 720 proteins by bond type (hydrogen bonds: 388 proteins) for the difference density at the site (grey) and for the charge from the joint fit (blue); bar and number: median. (b) C-C AUC of the joint fit against that of the difference density, one point per protein; the joint fit is higher in 601 proteins and lower in 119. (c, d) Median AUC over proteins by B of the site for C-C and C-N bonds, for the difference density, the matched filter on one site and the joint fit; bars are 95% bootstrap intervals of the median. Dotted lines: AUC 0.5 (`fig4_inverse`).

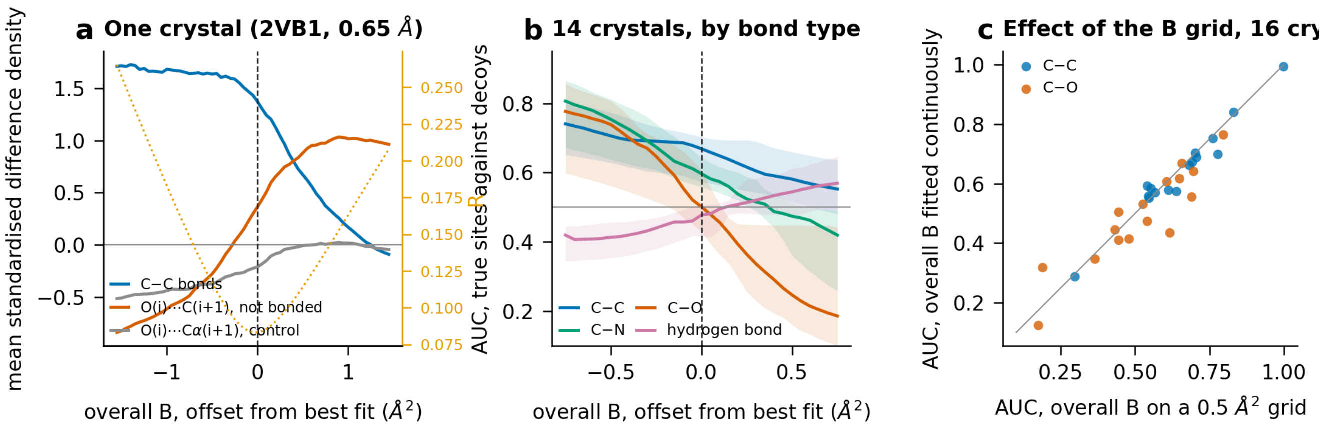


Figure S5. The overall B is part of the instrument. Sixteen crystals were scanned. (a) One crystal (2VB1, 0.65 A): mean standardised difference density at C-C bonds, at the non-bonded O(i)...C(i+1) contact and at a control pair, against the offset of the overall B from its best fit; dotted line and right axis: R. (b) AUC of true sites against decoys by bond type against the same offset, for the 14 crystals whose R-optimal B lies inside the range of this scan; line: median, band: quartiles. (c) AUC with the overall B on a 0.5 A^2 grid against the AUC with the overall B fitted continuously, all 16 crystals. The scan quoted in section SI-3 retains 15 of the same 16 (`fig5_overallB`).

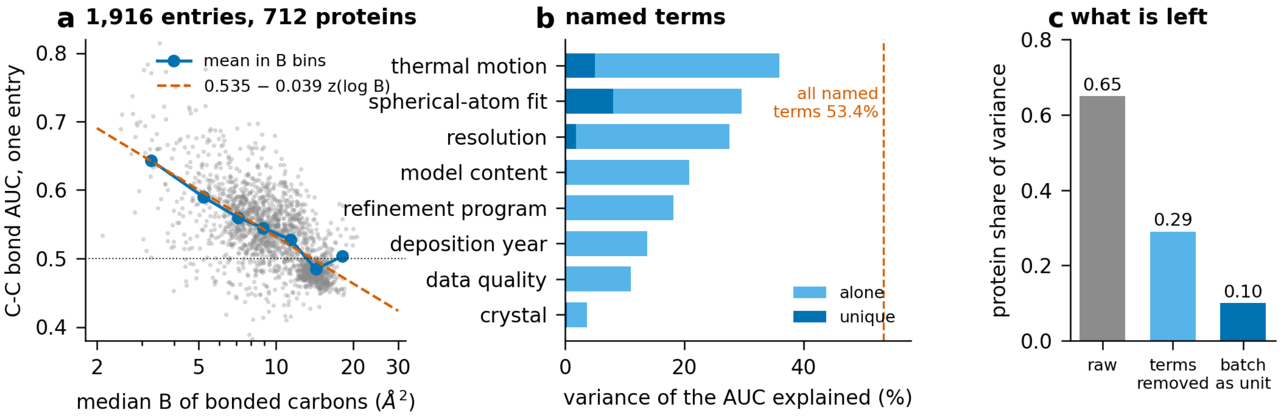


Figure S6. The order law. (a) C-C bond AUC of an entry against the median B of its bonded carbons (1,916 entries, 712 proteins; logarithmic B axis); blue: mean in bins of B; dashed red: AUC = 0.533 - 0.037 z(log B). (b) Variance of the AUC explained by each named block of entry-level quantities, alone (light) and uniquely (dark); dashed line: all named terms together, 54.1%. (c) Share of the variance that lies between proteins: raw (0.68), after the named terms are removed (0.40), and with the depositing batch as the unit (0.16) (`fig6_order_law`).

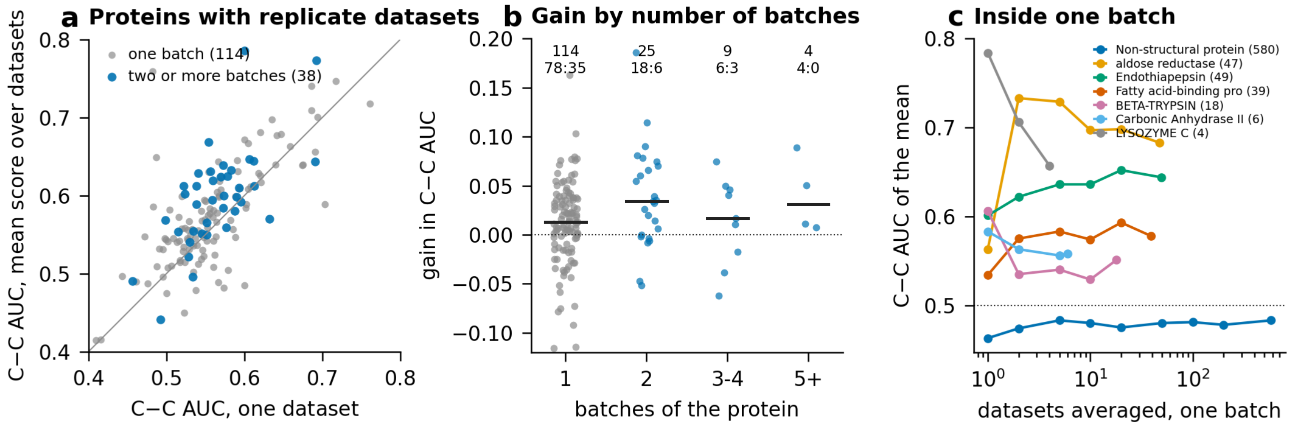


Figure S7. Several crystals of one protein. (a) C-C AUC of the mean score over all datasets of a protein against the AUC of one dataset, for proteins whose replicate datasets come from one depositing batch (114) and from two or more (38). (b) Gain in C-C AUC against the number of batches of the protein; over each group, the number of proteins and better : worse. (c) Inside one batch: C-C AUC of the mean of the first k datasets (logarithmic axis) for seven proteins, number of datasets in brackets (`fig11_multi_crystal`).

### Supplementary text

Supplementary sections SI-1 to SI-6 give in full the investigation summarised in sections 4.4 and 4.5 and at the end of section 4.10. They were computed on the models as deposited, before the hydrogens were placed (section 3.2), with the bond layer by element pair; the numbers are those of that condition.

#### SI-1. Density at the contact classes and the positional control

Ablating each class of the layer (R of the whole layer without the class minus R with it, over 1,996 entries, each protein counted once) shows the increment is carried by the broad hydrogen-bond classes: the n-to-pi* carbonyl class NPI (476 proteins helped, 39 hurt, sign p 1e-96), the hydrogen-bond classes HBX (475: 93) and HB (245: 67), and the water hydrogen-bond class HBW (374: 230). The side-chain aromatic and stacking classes (CHPI 159: 336, the stacking and cation-pi classes) and the all-atom anharmonic class lower R on net. Forcing all class widths into a narrow band removes the gain, because it collapses exactly these broad classes (the NPI contribution falls about sevenfold): the width of the hydrogen-bond cloud is where the signal lives, not noise to be regularised away. The proteins the layer makes worse are a tail of antibiotics and designed peptides on which the all-atom and ligand-ring classes overfit.

These ablation signs cannot yet be read as chemistry, and we report that as a result in its own right. The classes that help sit on the main chain, which carries a positive residual and a lower B; the classes that hurt sit on side chains, which carry the atom-centred negative well of section SI-4 and a higher B. The help/hurt split follows the main-chain against side-chain split exactly. To separate chemistry from position we trained, for every real class, a matched placebo class: one empty, non-bonded point per real site, matched on the two nearest-atom distances, the nearest modelled hydrogen, ln B, occupancy and the covalent-graph depth of its two nearest atoms, each placebo class carrying its own charge and width. The quantity that can be interpreted is the real-minus-placebo residual: the per-protein median of [R-drop(real class) minus R-drop(matched placebo)], positive when the real class lowers R more than its position-matched empty point.

| class | real beats placebo: not | sign p | real charge (two folds) | placebo charge |
|---|---|---|---|---|
| NPI (n-to-pi* carbonyl) | 521: 45 | 1e-104 | +2.31, +1.92 | +0.32, -0.01 |
| HBX (hydrogen bond) | 502: 110 | 1e-61 | +0.51, +0.25 | +0.04, +0.03 |

| class | real beats placebo: not | sign p | real charge (two folds) | placebo charge |
|---|---|---|---|---|
| HB (hydrogen bond) | 279: 76 | 2e-28 | +1.55, +0.62 | +0.13, +0.01 |
| AMPI (amide-pi) | 359: 116 | 6e-30 | +1.52, +0.26 | -0.08, -0.21 |
| SALT (salt bridge) | 243: 222 | 0.35 (null) | | |
| CHPI (C-H-pi) | 173: 365 | 8e-17 (worse) | | |

The hydrogen-bond and carbonyl family (NPI, HBX, HB, AMPI) survives the control: a Gaussian at the contact lowers R more than a Gaussian at a position-, B- and graph-depth-matched empty point, and its fitted charge is large where the placebo charge is near zero (NPI +2.3 against +0.3). This is real excess density at polar contacts, not the backbone-position artifact of the raw ablation. The salt-bridge, water-pi, cation-pi and anion-pi classes are null against their placebo (pure position), and the aromatic-stacking and C-H-pi classes are worse than their own position control. One caveat bounds the positive claim: the empty point is matched on position and B but not on the lone-pair and bond density that sits on the polar and carbonyl atoms themselves, and the inverse angular control (section SI-2) finds no specificity to the n-to-pi* donor-acceptor geometry. The surviving signal is therefore polar-contact and carbonyl-polarisation density beyond position, not detection of a named interaction by its geometry.

### SI-2. Inverse read-out on the models as deposited

On the models as deposited the question was put with decoys in place of matched points and with three scores. The answer depends on one setting of the instrument, the overall B (section SI-3), and on what the read-out responds to, which is the subject of section SI-4.

Sites are the midpoints of all atom pairs closer than the sum of their covalent radii plus 1.5 A. True bonds are positives; non-bonded pairs in that range are decoys. Three scores are compared: the IAM difference density at the site (raw, no parameter); a matched filter (single); and one charge per site refined jointly over all sites of the entry on the CEL basis (joint). The area under the ROC curve (AUC) is computed inside each entry, so that scale and data quality cancel. An AUC of 0.5 means that the score does not tell a bond from a decoy; a value below 0.5 means that it ranks them the wrong way round.

Over 720 proteins (figure S4a):

| type | raw | matched filter | joint | proteins above 0.5 (raw / joint) |
|---|---|---|---|---|
| C-C | 0.551 | 0.519 | 0.594 | 86% / 94% |
| C-N | 0.553 | 0.521 | 0.585 | 82% / 89% |
| C-O | 0.467 | 0.444 | 0.435 | 28% / 22% |
| hydrogen bond (388 proteins) | 0.425 | 0.431 | 0.448 | 18% / 12% |

The joint score is better than the raw density for C-C in 601 proteins and worse in 119 (figure S4b). The matched filter is worse than the raw density everywhere. A kappa base in place of the IAM lowers the C-C AUC in 594 of 681 proteins: the spherical valence term absorbs part of the bond signal.

The signal depends on the B of the site (figure S4c, d). C-C (joint) falls from 0.696 at B < 4 A^2 to 0.476 at B 17 to 25 A^2; C-O (raw) from 0.618 to 0.317. Restricted to the main chain, C-C and C-N stay above 0.5 at every B (C-C joint 0.730 to 0.551; tier values 0.643 and 0.618, above 0.5 in 94% and 89% of proteins). The values below 0.5 come from side chains and from the carbonyl oxygen. A loss of signal would end at 0.5; values below it are a signed effect, which section SI-4 measures directly and section SI-4.2 removes.

Hydrogen bonds are below 0.5 with every score. The positive residual found near hydrogen bonds in a scan of pair classes is the hydrogen atom itself: it vanishes when the hydrogen is in the model and shows no dependence on the donor-acceptor distance. The full CEL carries explicit hydrogen-bond and n-to-pi* classes, and we tested directly whether their forward gain reflects the interaction geometry or only the position of the site. It does not. At

O(i)...C(i+1) sites the standardised residual is +0.284 where the n-to-pi* geometric criterion is met and +0.201 where it is not, with no angular specificity: a positive residual of about +0.2 sits at any such main-chain midpoint, met or not. The same geometric control is therefore required for every non-covalent class before it is read as chemistry, which is the purpose of the placebo normalisation of section SI-1. A position-matched control (section SI-1) shows that the hydrogen-bond and carbonyl classes nonetheless carry density beyond position, while the aromatic, cation-pi and salt-bridge classes do not; but even the surviving family is not specific to the n-to-pi* donor-acceptor angle, so what CEL reads at these contacts is polar-contact density, not the named interaction geometry.

### SI-3. The overall B as part of the read-out

The first version of the pipeline fitted the overall B on a grid of 0.5 A^2. A reproducible positive residual between O(i) and C(i+1) of the main chain, which looked like direct density evidence for the n to pi* interaction, followed the grid step. A continuous scan on 15 entries showed that +0.25 A^2 of overall B moves the standardised residual at that site by +0.13 (in 15 of 15 entries) and the C-C AUC by -0.041 (in 14 of 15), and that at the R optimum the residual has no consistent sign. The signal was a scaling artefact. Figure S5a shows the scan for one crystal: as the overall B is raised the residual at the C-C bonds falls and the residual at the non-bonded O(i)...C(i+1) contact rises, and only R fixes the B at which either may be read. By bond type the sensitivity is ordered: per +0.25 A^2 the C-C, C-N and C-O scores fall by 0.028, 0.055 and 0.100 and the hydrogen-bond score rises by 0.047 (medians over 14 crystals; figure S5b). The more polar the bond, the closer its midpoint density lies to zero, and the more a small radial error decides its sign.

The overall B is now fitted continuously (two nested refinements, to 0.001 A^2) and the whole tier was recomputed. R falls in 1,898 of 1,927 entries. No tier value moves by more than 0.003, single entries move by 0.01 (median) and up to 0.06: the grid was noise at the level of the entry, not a bias (figure S5c). Two rules follow for any benchmark of this kind. The overall B must be fitted to the R optimum and reported. And it must not be tuned on the detection score: lowering B below the optimum raises the AUC monotonically (0.823 to 0.891 on 2VB1 while R goes from 0.083 to 0.265), because bond midpoints lie closer to atoms than decoy midpoints do.

### SI-4. The atom-centred term in the read-out

Most of what the inverse score responds to is not bonding density but a residual centred on the atoms. This section shows it at mobile sites (SI-4.1), measures it for every atom of the tier and removes it (SI-4.2), repeats the read-out outside the protein (SI-4.3) and on the non-covalent classes (SI-4.4), and asks whether the term follows from the distance environment of the atom (SI-4.5).

#### SI-4.1 At mobile sites the residual is centred on the atoms, not on the bonds

The inverse benchmark asks whether the residual density at the midpoint of a true covalent bond is higher than at decoy midpoints of the same entry. For ordered sites it is. For mobile sites the sign reverses. In the highest B class (B > 17 A^2) the standardised residual z is -1.15 at 0.55 A from the nearest atom centre, about -0.6 between 0.7 and 1.05 A, and returns to zero at 1.4 A, with no positive shell outside it (figure 4a). True bonds follow the same curve: at B 17 to 25 A^2 the median z of C-C, C-N and C-O bond midpoints is -0.42, -0.58 and -1.03. The feature is therefore centred on the atoms, not on the bonds. It is what a model leaves behind when a Gaussian of the refined width places more density at the atom centre than the crystal contains.

The consequence for bond detection is direct. Main-chain bonds stay above chance over the whole B range (C-C joint AUC 0.730 at B < 4 A^2, 0.551 at B 17 to 25 A^2). Side-chain C-N and C-O bonds fall below 0.5 (0.446 and 0.436), and C-O as a whole crosses 0.5 near B of 8 to 9 A^2 (figure 4d). A bond AUC below 0.5 is not evidence against bonding density. It is the negative well of the two bonded atoms overlapping at their midpoint; when the atom-centred term is solved and removed (section SI-4.2), the values below 0.5 disappear.

### SI-4.2 The atom-centred term, measured at the atoms and removed

The term can be measured where it is centred. For every atom of the tier (8.3 million) the IAM difference map was averaged on spheres of radius 0 to 1.6 A around the atom. In units of the standard deviation of the map of the entry, the value at the centre of a protein atom is +0.69 for B < 4 A^2, +0.54 at 6 to 8, +0.06 at 10 to 13, -0.34 at 13 to 17, -0.85 at 17 to 25 and -1.17 above 25 A^2, and at every B it falls to within about 0.2 of zero between 1.2 and 1.6 A (medians over 720 proteins). The atom-centred term is therefore not only a well. It is one term that changes sign near B of 10 to 13 A^2: an excess at ordered atoms and a deficit at mobile ones. Because the midpoint of a bond lies nearer to its atoms than the midpoint of a decoy does, the term raises the bond score where it is positive and lowers it where it is negative.

The term can be removed exactly rather than estimated. Writing the map as a sum of one radial function per atom of the crystal (17 nodes, to 1.6 A), the shell means of all atoms form a sparse linear system, because the mean of a neighbouring function over a sphere has a closed form. The system was solved in every entry (8.3 million atoms, 144 million atom pairs; planted functions are recovered with r of 0.997 or better), with no use of the labels and nothing transferred. The solved functions are narrower than the shell means: +0.70 at the centre for B < 4 A^2 and -0.93 above 25 A^2, and within 0.05 of zero from 1.1 A on for B below 10 A^2 (figure 5a). Subtracting the functions of all atoms within 1.7 A from the value at each midpoint gives a score without the atom-centred term. With it no bond type lies below 0.5 in any class of B (figure 5b): at B 17 to 25 A^2 C-C goes from 0.421 to 0.527, C-N from 0.381 to 0.543 and C-O from 0.309 to 0.531 (better in 506, 522 and 531 of about 550 proteins), and side-chain C-N and C-O go from 0.391 and 0.388 to 0.574 and 0.532 (better in 624 of 648 and 632 of 646 proteins). At B < 4 A^2 the same score is lower than the raw one for C-N (0.650 to 0.527) and C-O (0.627 to 0.523) and nearly unchanged for C-C (0.687 to 0.668).

What the term is, in electrons. In absolute units the value at the centre runs from +0.07 e A^-3 for atoms with B < 4 A^2 to -0.08 e A^-3 above 25 A^2 (the median standard deviation of a map of the tier is 0.093 e A^-3). Its integral over the sphere of 1.7 A is +0.10 to +0.12 electrons per atom below B of 10 A^2, passes through zero between 13 and 17 A^2, and is -0.20 electrons at 17 to 25 A^2 and -0.41 above 25 A^2. The integral decides the reading. An error in the width of the atomic Gaussian would move density between the centre and a surrounding shell and leave the integral near zero. What is found is a net excess at ordered atoms and a net deficit at mobile ones, with no compensating shell inside 1.7 A.

The two signs have different carriers. Below B of 6 A^2 the excess is largest on the backbone N and CA (+0.19 to +0.23 and +0.15 to +0.18 electrons), present on the carbonyl O (+0.16 to +0.20) and absent on the carbonyl C (+0.01). This is the ordering of the atomic charges of the peptide and of the atoms that carry hydrogen; it is the part of the density that a spherical valence term describes. Above B of 17 A^2 the deficit is largest on the side-chain O, N and S (-0.29 to -0.60 electrons at 17 to 25 A^2), smaller on the carbonyl O (-0.25) and on side-chain C (-0.15), and absent on the backbone N, CA and C (+0.06 to +0.08), which lose their excess only above 25 A^2 (figure 5e, f). The deficit follows the atoms that move most, not an element. We read the first sign as chemistry that the neutral spherical atom leaves out, and the second as density that a mobile atom does not have at its refined position. The data give the ordering; the reading is ours. Sulfur is the exception to both: its integral is already negative between B of 4 and 13 A^2 (-0.05 to -0.27 electrons), on about a hundred proteins.

The term is, to a good approximation, a fraction of the atom itself. The density of the IAM atom at the resolution of the entry is fixed by its element and its B, with no free number. One amplitude per atom times that density reproduces 83% of the variance of the solved functions, which have seventeen numbers per atom (median over 693 proteins, positive in every one). The amplitude is the fraction of the atom's own model density that is in excess or missing: +1.3 to +1.9% on the backbone N and CA below B of 10 A^2, passing through zero between B of 10 and 25 A^2 depending on the atom, -6% on side-chain N and O at 17 to 25 A^2, and -9 to -12% on side-chain atoms above 25 A^2, against -1 to -2% on the backbone N and CA there. No atom falls outside plus or minus 100%. B and the surroundings of an atom therefore do not add: B fixes the shape of what is missing, and the fraction that is missing is a second quantity, and the two together make up the atom. The remaining 17% is not negligible for the read-out. With the one-amplitude form subtracted in place of the solved functions, main-chain C-C reaches 0.634 instead of 0.696 and side-chain C-N 0.496 instead of 0.574: where the missing fraction goes, close to the atom, has a shape of its own that this form leaves out.

The two-component form states this directly and bounds it. Each atom is written as its model density plus a small charge term for its kind, and a fraction between 0 and 1 of the atom is taken out of the model position and placed in a shape common to atoms of the same kind and B class, so that the part in place and the part displaced add up to the whole atom. Shapes and charge terms were fitted on one half of the proteins and applied to the other, which leaves one number per atom. The displaced fraction has a median of zero below B of 10 A^2 and rises with B to 3% on the backbone N, 6.5% on the backbone O, 9.7% on side-chain C and 12.3% on side-chain N above 25 A^2. The charge term is +0.2 to +1.7% of the atom for every kind and agrees between the two halves. With one number per atom this form reproduces 59% of the variance of the solved functions out of fold (positive in 674 of 693 proteins), and the read-out over all sites is above the raw score for all three bond types: C-C 0.589 (better in 668 proteins, worse in 42), C-N 0.563 and C-O 0.530, where the free solution with seventeen numbers per atom gives 0.565, 0.525 and 0.518. It keeps main-chain C-N at 0.564, which the free solution takes to 0.500, and reaches 0.643 on main-chain C-C, 0.533 on side-chain C-N and 0.505 on side-chain C-O. Three limits are measured. Between B of 13 and 25 A^2 C-O stays at 0.47 and C-N at 0.50 to 0.51, where the free solution reaches 0.53 to 0.55. The shapes of the displaced part agree only weakly between the two halves (r = 0.29). And less than a fifth of the displaced electrons are found again within 1.7 A of the atom above B of 25 A^2. A spherical shape per kind of atom does not say in which direction the displaced part goes, and that is the quantity the next model has to carry.

Three readings follow. The values below 0.5 at mobile sites were the atom-centred deficit; without it they are gone. The dependence of the read-out on B (section SI-5.1) is, for C-N and C-O, the dependence of this term on B: after its removal both lie between 0.51 and 0.55 in every class of B, and only C-C keeps a gradient (0.668 to 0.508). And the term decides what a tier value means. Main-chain C-C becomes clearer without it (0.601 to 0.696, better in 629 of 677 proteins), whereas main-chain C-N falls from 0.587 to 0.500 (figure 5c): its raw read-out is not separable from the density centred on its atoms. Over all sites C-C moves from 0.555 to 0.565, C-N from 0.559 to 0.525 and C-O from 0.471 to 0.518. The score is conservative, because the spherical part of the bonding density itself is absorbed into the radial functions. The joint score of section SI-2 has not been recomputed without the term; the numbers of this section are for the raw density.

The term accounts for results elsewhere in this paper. It is why removing every property of the place of a site leaves the read-out unchanged (section SI-5.4): those properties shift the bonds and the decoys of a place together, whereas this term acts through the distance of a site from its atoms, and that distance is what separates a bond from a decoy. It is why the hydrogen-bond class reads above its controls in the raw density and below them without the term (section SI-4.4). And it is why the raw read-out falls with B: the excess that helps the bonds of ordered atoms and the deficit that hides the bonds of mobile ones are one function crossing zero.

The term is not a function of element and B alone. Forty class functions (five elements by eight classes of B), fitted on half of the proteins, agree between the halves (r = 0.91 for C, N and O), but applied to the other half they restore only part of the read-out at mobile sites (at B 17 to 25 A^2, C-C 0.504 and C-O 0.451). Inside an entry, log B explains 10% of the variance of the centre value over the protein atoms, and log B with the kind of atom 16% (medians over 1,881 entries); the rest is the individual atom, together with the noise of a solution that has seventeen numbers per atom. The term is a property of the individual atom, in line with the graph term of section SI-5.2; section SI-4.5 asks how much of it follows from the surroundings of the atom. A benchmark of bonding density should report the score with the atom-centred term removed next to the raw one.

#### SI-4.3 Residues that are not protein

The benchmark above is on protein atoms. The same structures hold 8,329 residues that are neither protein nor water and have more than one non-hydrogen atom (112,331 atoms): nucleic acids, cofactors, glycans, non-standard amino acids, bound small molecules and crystallisation additives. Nothing in the sites, the scores or the removal of the atom-centred term refers to the type of the residue, so these residues test the read-out outside the protein. Inside residues of 6 to 15 non-hydrogen atoms the raw read-out of all bond types together is 0.458 (400 proteins), below chance and below the protein value of the same tier (0.536). With the atom-centred term removed it is 0.544, against 0.534 for protein, and better than the raw score in 290 of 397 proteins. For residues of 16 atoms or more the two values are 0.519 and 0.548 (276 proteins), and for residues of 2 to 5 atoms 0.420 and 0.532 (201 proteins). The deficit of the raw score is carried by the mobile residues: at B above 17 A^2 the read-out goes from 0.394 to 0.526 and from 0.400 to 0.541 in the two larger classes. The bonds of non-protein residues are therefore read as well as those of the protein once the atom-centred term is removed, and they need the removal more (figure 5d). This is the read-out inside the residues; the contacts between a bound molecule and the protein are not separated here.

#### SI-4.4 Non-covalent classes, by who is in contact

The thirteen non-covalent classes were read against matched empty points (same distances to the two nearest atoms and to the nearest hydrogen, same B and occupancy; 2.48 million sites), raw and with the atom-centred term removed from sites and controls alike, and split by the residues of the two nearest atoms. Between protein atoms the carbonyl class NPI is above its controls in 516 of 622 proteins (median 0.581; 0.557 without the atom-centred term, 498 of 624) and the salt-bridge class in 358 of 446 (0.582; 0.571). The water hydrogen-bond class gains from the removal (0.515 to 0.555 between protein atoms, 0.537 to 0.566 at protein-water sites, above chance in 577 of 641 proteins). The hydrogen-bond class HB does the opposite: 0.548 raw (above chance in 271 of 354 proteins) and 0.452 once the term is removed (below chance in 277 of 354); its raw read-out is centred on the atoms. HBX is below its controls with both scores (0.425 and 0.437). The inverse read-out and the forward control of section SI-1 therefore do not rank the classes identically: NPI passes both, the salt bridge passes the inverse test only, and HB and HBX pass the forward control only. The two tests ask different questions, whether a site holds more density than a matched empty point, and whether a charge placed at the site lowers R more than a charge placed beside it. A class should be called supported by density only when it passes both.

Contacts between protein and non-protein residues are few in this tier: 8,538 sites to residues of six or more non-hydrogen atoms and 2,161 to smaller ones, against 1.08 million between protein atoms. Only two classes reach fifty proteins there, and neither differs from chance (HBX 0.478 raw and 0.488 with the term removed, 214 proteins; CHPI 0.531 and 0.525, 59 proteins). A class-by-class reading of the contacts of bound molecules needs a set of structures chosen for them.

#### SI-4.5 The term from the distance environment of the atom

If the term belongs to the atom, it should follow from where the atom sits. Each atom was described by its three-dimensional distance environment alone: the number of other atoms of the crystal, symmetry mates included, in twenty distance shells out to 8 A and six classes of neighbour (C, N, O, S and other, H, water O). These are 120 numbers, with no density, no B and no label in them. A linear model of the solved radial function on these numbers, one set per kind of atom, was fitted on half of the proteins and applied to the other half, next to the same model with B in place of the environment, and with both.

Out of fold and inside an entry, the kind of atom alone explains 1.4% of the variance of the centre value over the protein atoms, the kind with B 9.0%, the kind with the distance environment 9.6%, and all three 12.0% (2.4, 9.0, 10.9 and 13.3% for the whole function; medians over 693 proteins). Geometry without B does as well as B, and adds to it in 553 of 693 proteins. The shares are small, as expected of a linear, additive form set against a solution with seventeen numbers per atom.

The test that matters is the read-out with the predicted functions subtracted in place of the solved ones. For main-chain C-C the raw AUC is 0.601 and the solved term gives 0.696; the prediction from B gives 0.606 and the prediction from the distance environment 0.639, better than raw in 604 of 672 proteins. For side-chain C-N the four values are 0.391, 0.574, 0.455 and 0.527 (634 of 648 proteins), and for side-chain C-O 0.388, 0.532, 0.452 and 0.510 (640 of 645). The environment alone therefore recovers about 40% of what the solved term recovers for main-chain C-C and three quarters or more for the side-chain bonds, in proteins that took no part in the fit. B does better only at the mobile end (C-C at B above 25 A^2: 0.513 from B, 0.455 from the environment, 0.508 solved), and the environment at the ordered end, where the prediction from B lowers the read-out (C-C at B < 4 A^2: 0.607 from B, 0.666 from the environment, 0.668 solved). The coefficients of the individual shells are not stable between the halves (r = 0.25), because neighbouring shells are collinear: the predictions transfer, the single numbers should not be read. Predicting instead the fraction of the two-component form of section SI-4.2, with the shape fixed by B, does no better: the environment explains 8.4% of the variance of the solved functions, B 8.0% and both 10.4%, and main-chain C-C reaches 0.611. Predicting the bounded displaced fraction of the two-component form, by a logistic function of the environment, of B or of both, explains 0.9%, 1.5% and 4.1% of the variance of the solved functions and gives main-chain C-C 0.617, 0.616 and 0.622 and side-chain C-N 0.471, 0.465 and 0.474. The additive prediction above (0.639 and 0.527) remains the best out-of-fold predictor; the displaced fraction of an atom is measured here and is not yet predicted from the structure.

The atom-centred term is thus set in part by the three-dimensional arrangement of the atoms around it, and that part can be computed for a structure, or for an arrangement of atoms that has not been observed, before any density is measured.

### SI-5. Determinants of the atom-centred term

The term of section SI-4 is positive where atoms are well ordered and negative where they are mobile. This section asks what sets it: the order of the entry and of the site (SI-5.1), the position of the atom in the covalent graph (SI-5.2), and the records of disorder in the deposited model (SI-5.3). Section SI-5.4 lists the corrections by named terms that do not remove it.

#### SI-5.1 Order: B, packing and rigid-body motion

Between entries. 68% of the variance of the C-C AUC lies between proteins. Eight named blocks of entry-level quantities explain 54.1% together (figures 4b and S4a): thermal motion 36.4% alone, fit of the spherical-atom model 30.6%, resolution 26.5%, model content 19.9%, refinement program 17.3%, deposition year 14.7%, data quality 12.7%, crystal 4.3%. Their unique parts are small (thermal motion 5.0%, spherical-atom fit 8.5%, all others below 2%): the blocks measure the same thing. One explicit law carries two thirds of the explained part (figure S6a):

$$\mathrm{AUC} = 0.533 - 0.037\, z(\log B_{\mathrm{CC}}), \qquad R^2 = 36.2\%$$

with $B_{\mathrm{CC}}$ the mean B of the bonded carbon atoms of the entry, standardised over the tier. After all named terms are removed, 40% of the remaining variance is still between proteins, 16% when the depositing batch is the unit (figures 4c and S4b).

Between sites of one entry. Two explicit terms account for the B of a site. Packing: the logarithm of the number of atoms around the site explains 33.1% of the variance of log B. From the loosest to the densest of five packing classes (75 to 126 atoms within 8 A) the mean B of the site falls from 12.4 to 7.3 A^2 and the C-C AUC rises from 0.493 to 0.607 (674 proteins; figure S1a, b). The same bond in two crystals is the direct test: for 4,365 bonds seen in two crystal forms of the same protein, the copy in a lattice contact has the lower B in 66.7% of cases (ratio 0.89) (figure

S1c). Rigid-body motion: the B field predicted from the deposited anisotropic tensors by a rigid-body fit per chain (Schomaker & Trueblood, 1968; as parameterised by TLS, Winn et al., 2001) correlates with the observed B with r = +0.69, against +0.03 for a control with the directions randomised (figure S2a), and explains 49.3% of the variance of log B (figure S2c). Together the two terms explain 56.0% (rigid body unique 22.0%, packing unique 6.7%, shared 27.3%), and 44.0% is not explained (figures S2c and S4c). The observed amplitude is 2.36 times the rigid-body prediction (figure S2b), so the rigid-body term is a lower bound of the correlated motion.

Order acts on the read-out largely through one term. Section SI-4.2 shows that the residual centred on the atoms changes sign with B and that, once it is removed from the raw score, the read-out of C-N and C-O no longer depends on B. What this section establishes is therefore what sets the size and the sign of that term.

#### SI-5.2 At equal B, the residual depends on the position of the atom in the covalent graph

B does not account for all of this. At equal B, z falls with the graph distance of the atom from the backbone and falls again for terminal atoms (figure 4b):

$$z = 0.443 - 0.610\,\log(B/B_0) - 0.155\,\mathrm{depth} - 0.255\,\mathrm{term}$$

where depth is the number of bonds from the outer atom of the bond to the nearest backbone atom (N, CA, C; capped at 5) and term is 1 if the bond involves an atom of degree one. These four numbers explain 9.15% of the site-level variance, against 9.69% for fourteen separately fitted chemical classes, and they halve the spread of the class residuals (0.292 to 0.137; figure 4c). The coefficients are the same on two disjoint halves of the proteins. The description is explicit and uses only the covalent graph, so it applies to any residue or ligand without a per-molecule calculation.

Chemistry-specific explanations account for little of it. Carboxylates, the groups most exposed to radiation damage, lie only 0.1 to 0.25 below amides at the same graph position. The dominant variable is the position, not the functional group.

#### SI-5.3 A ladder of disorder: B, alternative conformations, partial occupancy, missing atoms

If the graph term is mobility that B does not capture, the other records of mobility in the deposited models should rise along the same axis. They do. Over 165 atom positions of the twenty amino acids, from depth 0 to depth 4 the median B rises from 9.3 to 14.7 A^2 (x1.6), the fraction of atoms modelled with alternative conformations from 7.9 to 13.1%, the fraction with partial occupancy from 0.7 to 2.5%, and the fraction of atoms absent from the model from 0.02 to 0.87% (x48). Over the same range z at equal B falls from +0.31 to -0.46. Across atom positions the rank correlation of z at equal B is -0.81 with depth, -0.71 with B, -0.63 with the missing fraction, -0.46 with partial occupancy and -0.25 with alternative conformations (figure S3a to c).

The four records are steps of one ladder (figure S3d). Small displacements are absorbed by B. Larger ones are modelled as discrete alternatives. Beyond that the atom is given partial occupancy, and finally it is left out. The negative well measured here sits between the first and the second step: motion too large or too anharmonic for one Gaussian, but not resolved into separate positions by the crystallographer.

That side chains are more mobile than the main chain, that B factors underestimate the true heterogeneity, and that many side chains have unmodelled alternative conformations is established (Schneider et al., 2014; Kuzmanic, Pannu & Zagrovic, 2014; Lang et al., 2010; van den Bedem et al., 2009). What is added here is a measurement at equal B over a whole resolution tier (1,916 entries, 712 proteins), expressed in four numbers on the covalent graph. To our knowledge this has not been reported.

#### SI-5.4 Limits of a correction by named terms

Three direct uses of the graph term were tested and all failed. Subtracting it from the inverse scores does not raise the AUC. The deposited anisotropic displacement tensors do not carry it. Scaling B or occupancy by depth in the forward model raises R, because the deposited values already sit at the R optimum of a Gaussian model. The deficit is therefore not a mis-set parameter of the Gaussian model. It is the shape that a Gaussian cannot take. Nor does removing every label-free term together: with B, graph position, packing, lattice contact and the rigid-body B fitted on half of the proteins and taken out of the site scores of the other half, no tier value moves by more than 0.02 (C-C 0.551 to 0.549, C-N 0.552 to 0.554, C-O 0.469 to 0.476; 710 proteins). These terms describe the place of a site and move the bonds and the decoys of that place together.

### SI-6. Several crystals of one protein as the unit of replication

The last property of the instrument is its unit of replication: what has to be repeated for the read-out to improve.

For 152 proteins the tier holds more than one dataset. Averaging the raw score over the datasets of a protein raises the C-C AUC from 0.551 to 0.562 when all datasets come from one depositing batch (114 proteins, better in 78, worse in 35, $p = 6e\text{-}5$) and from 0.559 to 0.599 when they come from two or more batches (38 proteins, 28: 9, $p = 0.003$) (figure S7a, b). A three-level model (site, batch, dataset) gives a dataset noise of 0.807 and a batch term of 0.379 in units of the standardised score: datasets deposited together share their atomic model and their errors. Inside one batch the average saturates or falls with the number of datasets (figure S7c; beta-trypsin 0.606 to 0.551 over 18, lysozyme 0.784 to 0.657 over 4). The unit of replication of a density benchmark is the independent deposition, not the dataset.

The gain does not depend on what is removed from the scores first. With B, graph position, packing, lattice contact and the rigid-body B taken out of every site score (section SI-5.4), the same averaging gives 0.548 to 0.557 inside one batch and 0.557 to 0.596 across batches (29: 8), against 0.551 to 0.562 and 0.559 to 0.599 without the removal. Normalising the sites and then averaging ends where averaging alone does.

The posterior bond probability of the same model also rises for proteins with a single dataset (0.557 to 0.588, 541 proteins). That part comes from the prior on site B and bond type and is not an effect of averaging; the numbers above are the averaging effect alone.

### SI-7. Scans for unexplained density

Because the benchmark standardises the residual inside each entry, it can be turned on atom pairs that are not bonded. Two scans were run on the whole tier: 740 classes of non-bonded pairs defined by residue and atom names, and a second scan by generic chemistry that includes metals, ligands and modified residues. The strongest positive classes are hydrogen bonds (the unmodelled hydrogen, section SI-2) and coordination bonds to metals (Fe-S +1.25, Fe-N +1.07, Cys S-Zn +0.60). The most negative are the centres of aromatic rings (Phe -0.45, in 85% of 530 proteins) and bonds between electron-rich atoms (S-O -1.17, C-Cl -1.09, N-O -0.92, S-S -0.68).

Disulfides are the case a reader of a bonding-density paper asks for first, and they are instructive because they fail the naive version of the test. Across the tier the Cys S...S midpoint of 1,249 disulfides in 130 proteins carries a negative standardised residual (mean -0.68, positive in only 32 percent of proteins), not the positive peak that C-C and C-N bonds show. This is not a defect of the model. The difference density at the midpoint of a bond between two electron-rich atoms is negative in small-molecule charge-density work, and the electron-rich list here (S-O -1.17, N-O -0.92, C-Cl -1.09, S-S -0.68) is negative throughout. A single-structure inverse test can nonetheless appear to recover the disulfides of a protein, because the true S...S pairs sit at about 2.05 A while the decoys lie further out, and a midpoint closer to its two flanking atoms carries more density whatever the bonding (section SI-3). Ranking true disulfides above distant decoys therefore measures proximity, not bonding density, and it is exactly the confound that the within-entry standardisation and the position-matched controls of this paper are built to remove. Read correctly, the disulfide midpoint is a negative, and an honest one.

One lead was followed to the end (iron-sulfur clusters have been a target of charge-density analysis at ultra-high resolution, Hirano et al., 2016). The Fe...Fe midpoints of iron-sulfur clusters are positive in 20 of 21 crystals, and the density is higher toward the neighbouring Fe than in free directions (19 of 21) or between two sulfur ligands that have no Fe behind them (20 of 21). It is nevertheless not bonding density. Every Fe carries an isotropic positive shell at 1.2 to 1.5 A, and the sum of the shells of the two atoms reproduces the profile along the Fe...Fe line at every radius with no free parameter (midpoint: observed +2.20, predicted +2.18; excess positive in 8 of 21 entries, $p = 0.76$). The shell is a property of how Fe is modelled (candidates: scattering factor of the neutral atom against the ion, anharmonic displacement, series termination) and was not pursued.

No density feature was found at pair midpoints that calls for a new kind of interaction. Under the matched position control (section SI-1) the hydrogen-bond and carbonyl classes do carry density beyond position, but without specificity to the named interaction geometry (section SI-2), while the salt-bridge, cation-pi and stacking classes carry none. We report the scans because the negative result has the same coverage as the positive ones: the whole tier, every pair class.
